# N-acetylcarnosine attenuates age-associated declines in multi-organ systems to improve survival

**DOI:** 10.1101/2025.08.19.671148

**Authors:** Edwin R. Miranda, Justin L. Shahtout, Shinya Watanabe, Jan Spaas, Norah Y. Milam, Grace Neiswanger, Benjamin Werbner, Deborah Stuart, Sohom Mookherjee, Jack Wilson, Michael Judge, Isabella Y. Goh, Tyler Slater, Molly R. Gallop, Guoli Hu, Takuya Karasawa, Jillian K. Landers, Bryann Rainbow, Kyle T. O’Connor, Nicholas J. Black, He Lan, Linda S. Nikolova, Ying Li, Crystal F. Davey, James E. Cox, Sihem Boudina, Courtney M. Karner, Natalia S. Harasymowicz, Nirupama Ramkumar, J. David Symons, Amandine Chaix, Jonathan Z. Long, Micah J. Drummond, Katsuhiko Funai

## Abstract

Histidine containing dipeptides (HCDs) such as N-acetylcarnosine are endogenous metabolites that are ergogenic and mitigate metabolic dysfunction. We previously demonstrated that short-term N-acetylcarnosine treatment is highly efficacious in protecting muscle atrophy induced by disuse. Here we demonstrate that a 6-months treatment of N-acetylcarnosine attenuates a broad spectrum of age-associated maladies and improved survival by ∼50% in female mice. A comprehensive survey of organ systems revealed that N-acetylcarnosine prevents decline in adiposity, diastolic function, vasodilation, muscle strength, and bone density. Together, N-acetylcarnosine substantially delays the onset of system-wide end-stage pathology to prolong lifespan. As an endogenously present metabolite, treatment with N-acetylcarnosine may be a safe and promising intervention to promote healthy aging in humans.

## Introduction

With the aging global population, there remains an unmet need for therapies to mitigate age-related pathologies. While lifestyle modifications such as caloric restriction^1,2^ and exercise^3^ are effective strategies for prolonging health during aging, no equivocal pharmacologic strategies exist. Histidine containing dipeptides (HCDs) are endogenously produced metabolites primarily in excitable tissues^4^ such as skeletal and cardiac muscle by conjugating histidine with beta-alanine or gamma-aminobutyric acid (GABA) in the case of homocarnosine. The supplementation of β-alanine, which is the rate limiting metabolite for HCD production, is ergogenic in humans, improving aerobic performance and muscle force production^5^. Conversely, decline in the HCD N-acetylcarnosine predicts mental and physical frailty in octogenarians^6^. The purported primary mechanism for these health benefits involves the scavenging of reactive carbonyl species, especially lipid-derived carbonyls such as 4-hydroxynonenal (4HNE) via the imidazole ring of histidine^7^. 4HNE and other reactive carbonyl species are generated via oxidation of their metabolic precursors. For example, 4HNE is produced primarily via peroxidation of ω6- polyunsaturated lipids. Formation of reactive carbonyls are accelerated in contexts of oxidative stress such as aging^8^. These reactive carbonyls non-enzymatically and irreversibly bind to proteins on nucleophilic residues incurring consequences such as impaired protein structure, function, and turnover^9,10^.

HCD supplementation mitigates impairments in cardiovascular, musculoskeletal, neuronal and metabolic systems in contexts other than aging^11^. For example, supplementation with carnosine or its analog carnosinol improved glucose tolerance in mice and humans with obesity^12,13^. We recently demonstrated the ability of carnosine or N-acetylcarnosine supplementation to attenuate accumulation of 4HNE-modified proteins and preserve muscle mass and *ex vivo* force production in mice subjected to 7-days of inactivity^14^. Similarly, increasing muscle carnosine in older adults by supplementing with β-alanine improved muscle power and work capacity^15^. Nonetheless, there has never been a study that examined the efficacy of long-term HCD supplementation on aging *per se*. HCDs have excellent safety profiles with only mild side effects^13,15,16^, making them attractive to administer to humans over the course of aging. Herein, we performed a 6-month preclinical N-acetylcarnosine trial in 18-24 months old mice and performed systematic phenotyping of their organ systems to determine its effects on age-related health decline. We chose N-acetylcarnosine primarily because we believed it would be more a translatable therapy for humans due to its superior stability in human plasma compared to carnosine. Unlike mice, humans ubiquitously express carnosinase (CN1) which hydrolyzes carnosine and other HCDs limiting their stability in circulation. However, N-acetylcarnosine, which is largely unique to humans, is not susceptible to CN1 hydrolysis.

C57Bl/6 mice from the aged rodent colony at the National Institute on Aging were acquired at 17 months of age. Following acclimation in our facility, the intervention was initiated at 18 months of age and continued for 6 months. During the final month of the intervention, *in vivo* physiological metrics spanning metabolic, neuro-endocrine, cardiovascular, renal, and musculoskeletal systems were assessed. After the 6-month intervention, mice were anesthetized, and tissues were collected for various assessments (Figure 1A).

**Figure 1.**
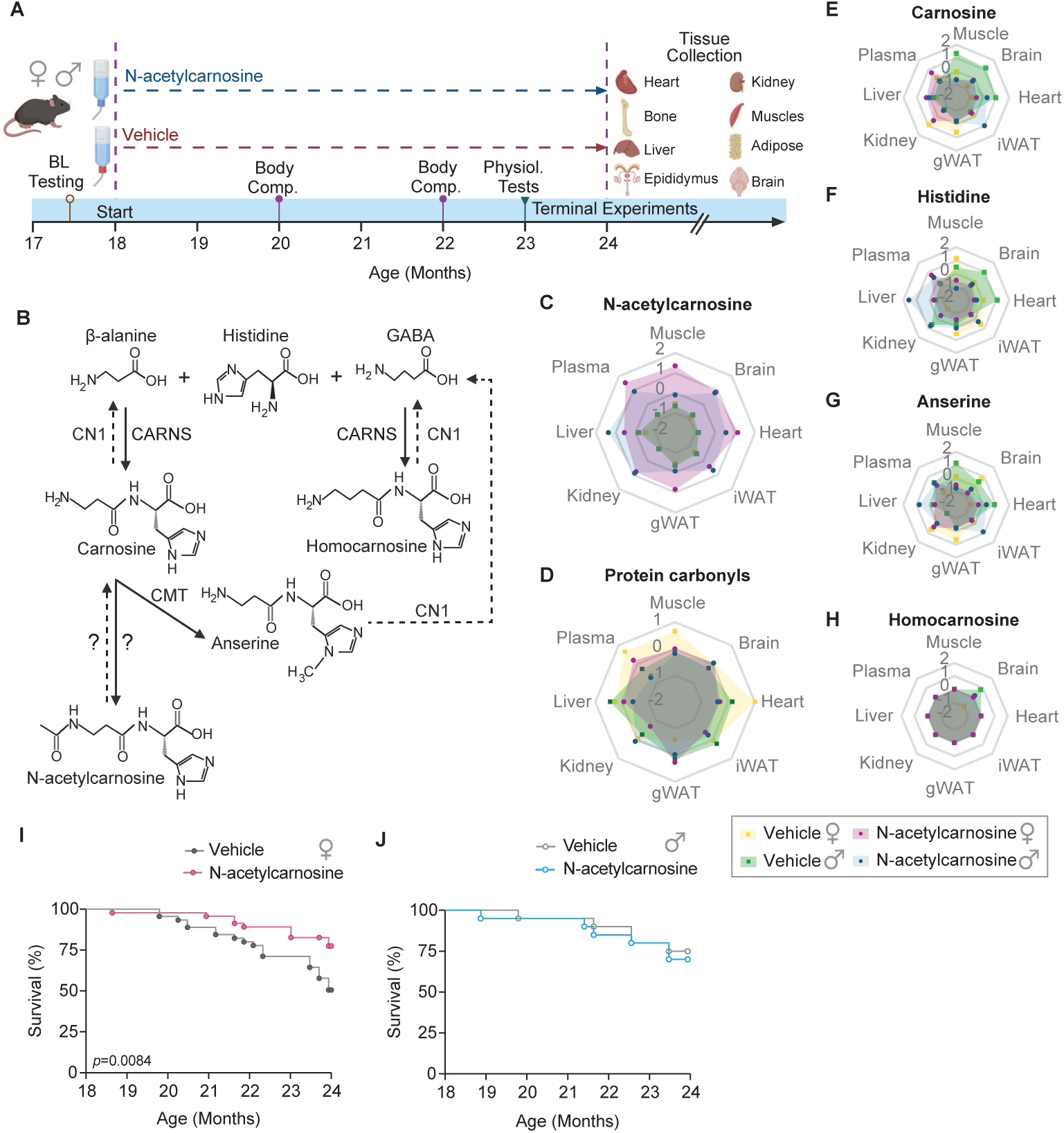
N-acetylcarnosine supplementation preferentially improves female mouse survival. **A)** Schematic of the preclinical trial design. C57Bl/6 mice from the NIA aging colony were acquired at 17 months of age. After acclimation, mice underwent baseline testing before being randomized to receive either vehicle or N-acetylcarnosine-supplemented water for 6 months. Mice were weekly monitored for body weight and food and water consumption. Various physiologic tests were conducted during the 5^th^ month of the intervention. Mice were euthanized and tissues were harvested after 6 months of intervention period. **B)** Schematic representation of HCD metabolism. Radar plots of **C)** N-acetylcarnosine, **D)** protein carbonyls, **E)** Carnosine, **F)** Histidine, **G)** Anserine, and **H)** Homocarnosine demonstrate enrichment of N-acetylcarnosine. Values in radar plots are Z-scores. Corresponding concentrations and statistical analyses are in **Supplemental Figure 1**. Vehicle treated female **(I)** and male **(J)** mice experienced typical survival rates for this strain and colony (approximately 50% and 75% respectively) at 24 months. Female mice (starting n=45 vehicle, n=46 N-acetylcarnosine) treated with N-acetylcarnosine improved their survival probability by ∼50%. Statistical testing for HCD enrichment was performed via two-way repeated measures ANOVA with Bonferroni post-hoc test where appropriate. Survival was assessed via generating Kaplan-Meier survival curves and testing the disparity of the curves via Mantel-Cox test. Comparisons were considered statistically significant if *p*<0.05. HCD; Histidine Containing Dipeptide, GABA; gamma-aminobutyric acid, CN1; carnosinase 1, CARNS; carnosine synthase. Schematic images were generated via BioRender.

## Results

To confirm efficacy of our treatment, N-acetylcarnosine and other histidine-related metabolites (Figure 1B) were quantified in various tissues. As anticipated^4^, N-acetylcarnosine was only appreciably quantifiable in tissues from N-acetylcarnosine-treated mice (Figure 1C, Supp. Figure 1). Given the role of N-acetylcarnosine to scavenge free carbonyls and prevent protein carbonylation, we also measured protein carbonyls in these tissues. Consistent with this notion, abundances for protein carbonyls were lower in several tissues from N-acetylcarnosine-treated mice compared to vehicle-treated mice (Figure 1D, Supp. Figure 1). Carnosine, and the other HCDs assayed were largely not enriched and some were even reduced by N-acetylcarnosine treatment (Figure 1E-H, Supp. Figure 1). In skeletal muscle, heart, and gonadal white adipose tissue (gWAT), N-acetylcarnosine enrichment was greater in female mice than in male mice. This observation is in accord with many of the sex-specific effects we observed on physiologic functions assayed.

At the 24-month timepoint, N-acetylcarnosine increased survival by ∼50% only in female mice (Figure 1I&J). In line with published female survival rates for this colony^17^, approximately half of the vehicle-treated female mice survived to approximately 24 months of age, while more than 75% of N-acetylcarnosine-treated female mice survived to this age. N-acetylcarnosine did not influence survival in male mice by the end of the study (∼24-month of age) (Figure 1J). It is important to note that 75% survival for 24-month old male mice in this colony is in line with published survival rates for male mice in this colony^17^. Concomitant with the increased survival, N-acetylcarnosine promoted a retention of body weight that was also specific to female mice (Figure 2A&B). The effect of N-acetylcarnosine on maintaining body weight was not explained by differences in lean mass (Figure 2C&D) but instead was exclusively accounted for by its ability to retain fat mass (Figure 2E&F). Importantly, these sex-specific effects were not explained by differences in food consumption, or exposure to N-acetylcarnosine via water consumption (Supp. Fig 2A-F).

**Figure 2.**
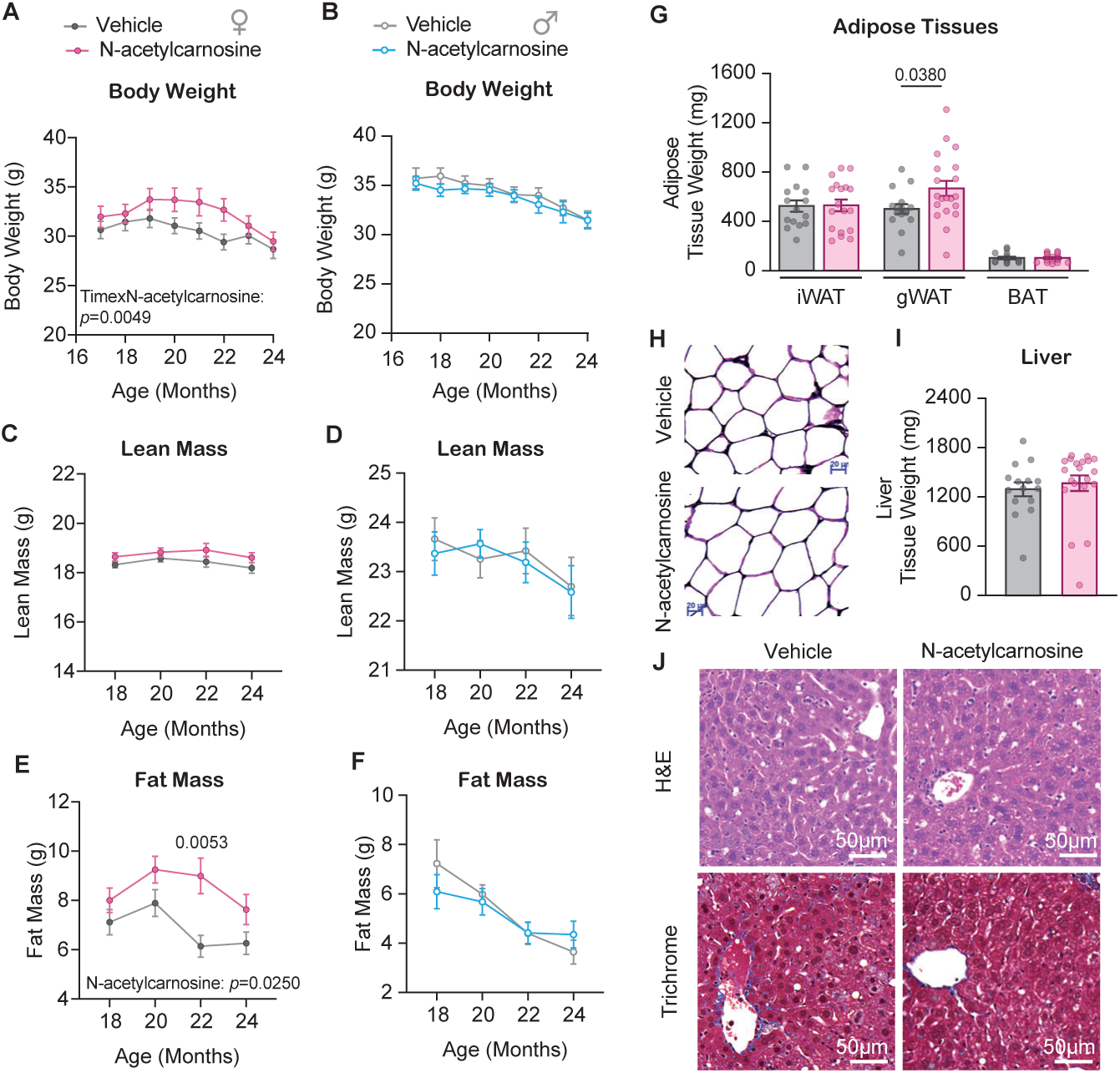
N-acetylcarnosine supplementation preserves body mass in female mice. Body weights in the female (starting n=39 vehicle, n=40 N-acetylcarnosine) **(A)** and male (starting n=20 per group) **(B)** mice fell precipitously over the course of the trial regardless of treatment. Female mice treated with N-acetylcarnosine were able to delay the decrease in bodyweight. Bodyweights are reported as one-month averages and are presented as Mean ± SEM. N-acetylcarnosine did not affect lean mass in female **(C)** or male **(D)** mice. N-acetylcarnosine preserved fat mass in female mice **(E)** (n=57 per group) but not male **(F)** (n=20 per group) mice. **G)** Preservation of fat mass in female mice was attributed to larger gWAT not iWAT or BAT fat pad (n=16 vehicle, n=20 N-acetylcarnosine). **H)** H&E stain of gWAT tissues did not reveal differences in adipocyte size between groups. Despite increased adiposity, liver weight **(I)** was not altered by N-acetylcarnosine (n=15 vehicle, n=20 N-acetylcarnosine. **J)** H&E and trichrome stains did not indicate altered steatosis or fibrosis in livers. Statistical testing for bodyweight and body composition comparisons was performed via one-way repeated measures ANOVA with Bonferroni post-hoc test where appropriate. Statistical testing for comparison of tissue weights was performed via unpaired student t-test. Comparisons were considered statistically significant if *p*<0.05. gonadal white adipose tissue; gWAT, inguinal white adipose tissue; iWAT, Brown Adipose Tissue; BAT.

Short-term N-acetylcarnosine or other HCDs may mitigate metabolic dysfunction associated with obesity^12,18–22^. Consistent with the body composition data, N-acetylcarnosine treatment promoted greater gWAT mass without influencing inguinal white adipose tissue (iWAT) or brown adipose tissue BAT mass in female (Figure 2G&H) but not male (Supp. Figure 3A) mice. It’s noteworthy that N-acetylcarnosine was robustly enriched in gWAT only in female mice (and not in male mice) whereas it was equally enriched in iWAT between sexes (Figure 1B, Supp. Figure 1). However, protein carbonyls were equivocal in between groups in male and female mice (Figure 1C, Supp. Figure 1). Similarly, liver, another metabolic organ influenced by adiposity, did not appear to be affected in mass or histology by N-acetylcarnosine (Figure 2I&J, Supp. Figure 3A&B). Systemic glucose tolerance was also not different between groups (Figure 3A, Supp. Figure 3C,D,K).

**Figure 3.**
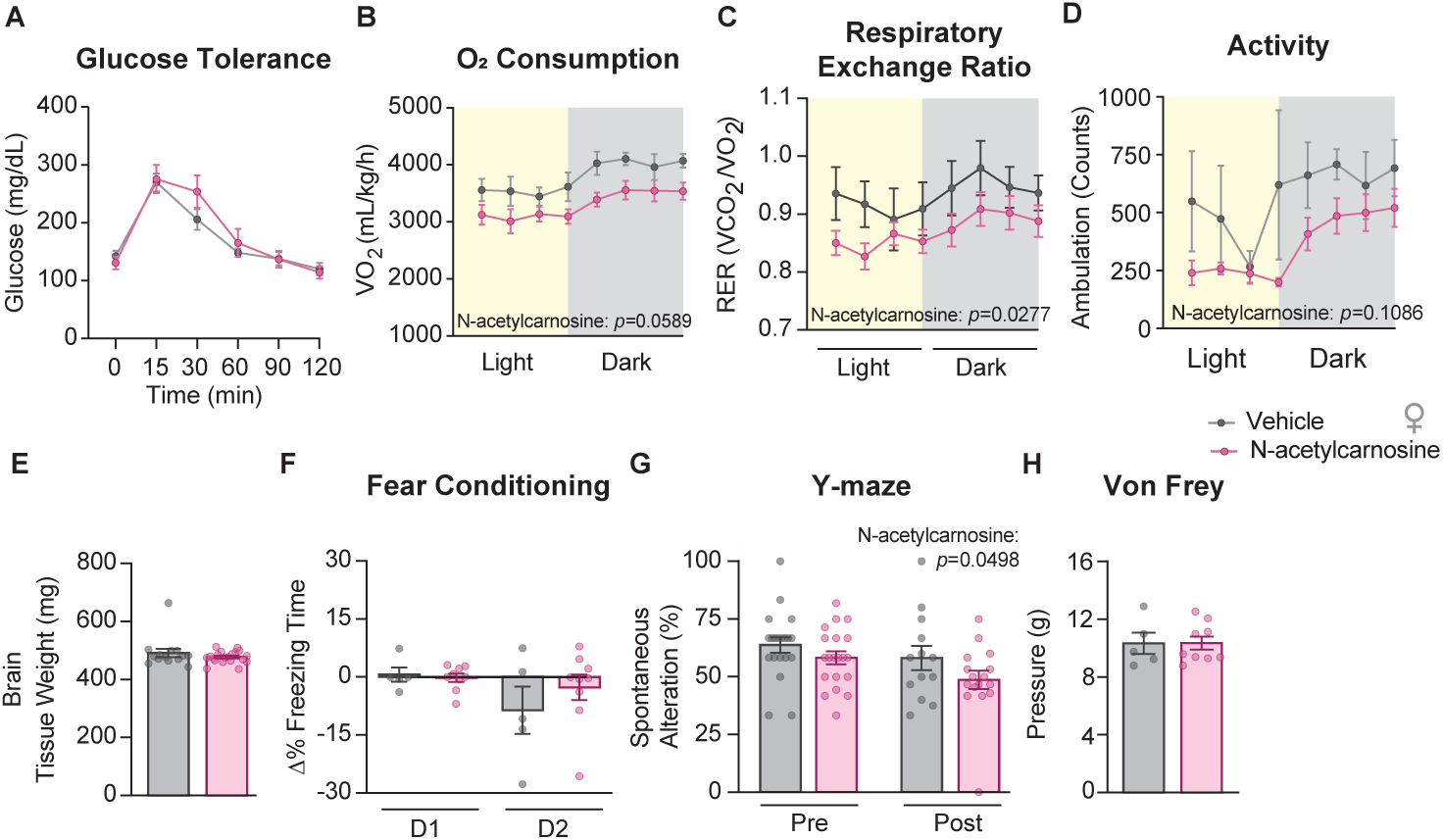
N-acetylcarnosine supplementation does not alter metabolic or neuroendocrine system in female mice. **A)** Glucose tolerance via intraperitoneal glucose tolerance test (IPGTT) was not different between treatment and vehicle mice (n=6 vehicle, n=9 N-acetylcarnosine). 72-hour indirect calorimetry revealed suppressed energy expenditure (VO_2_) **(B)** and respiratory exchange ratio (RER) **(C)** during both light and dark phases (n=5 vehicle, n=8 N-acetylcarnosine). Respiratory exchange ratio is the volume of CO_2_ consumed (VCO_2_) proportional to the O_2_ consumed (VO_2_) and indicates substrate utilization whereby a value closer to 0.75 indicates complete reliance on fat oxidation and a value closer to 1 indicates complete reliance on glucose oxidation. Ambulatory frequency was also suppressed in N-acetylcarnosine mice **(D)** (n=5 vehicle, n=8 N-acetylcarnosine). **E)** Brain weights were not different between treatment groups (n=14 vehicle, n=19 N-acetylcarnosine). **F)** Change in time spent freezing before and after an audible tone that proceeded with a shock on the first (condition day) but not the second (follow up) day was not different between groups indicating no effect of N-acetylcarnosine on conditional memory formation (n=5 vehicle, n=9 N-acetylcarnosine). **G)** Frequency of novel, non-consecutive exploration (alterations) between arms in the Y-maze was not different between groups indicating no effect of N-acetylcarnosine on short-term memory (n=20/14 pre/post vehicle, n=20/16 pre/post N-acetylcarnosine). **H)** Peripheral pain tolerance assessed via Von Frey hair filaments were not different between groups (n=5 vehicle, n=9 N-acetylcarnosine). Data are represented as Mean ± SEM. Statistical significance was assessed via two-way repeated measures ANOVA with Bonferroni correction, or unpaired t-test where appropriate. Comparisons were considered statistically significant if *p<*0.05. RER; respiratory exchange ratio, CLAMS; comprehensive lab animal monitoring system.

We further examined the effect of N-acetylcarnosine on systemic metabolism by Indirect calorimetry. Surprisingly, whole-body O_2_ consumption and energy expenditure were consistently reduced in the N-acetylcarnosine treatment group in both sexes (Figure 3B, Supp. Figure 3E,F,L), despite greater body weight in N-acetylcarnosine-treated female mice (Figure 2A). Respiratory exchange ratio (RER), defined by CO_2_ produced per O_2_ consumed, is an indirect parameter to estimate systemic substrate preference. An RER of 1.0 indicates exclusive glycolytic oxidation, while a RER of 0.7 indicates exclusive fat oxidation. N-acetylcarnosine treatment did not have any effect on RER in male mice (Supp. Figure 3H). Somewhat surprisingly, N-acetylcarnosine treatment reduced RER in female mice (Figure 3C), normally a signature of improved metabolic health^23^. These effects on metabolism are most likely explained by reduced ambulatory activity that was observed in the female (Figure 3D, Supp. Fig 3M&N) but not male (Supp. Figure 3G,I,J) N-acetylcarnosine-treated mice. Overall, N-acetylcarnosine preserved fat mass in female mice without signs of metabolic dysfunction. In the context of aging, retention of adiposity represents the ability of animals to maintain energy reserve which may delay frailty and promote longevity.

For male mice, we also assessed the effect of N-acetylcarnosine on fertility by assessing sperm count and sperm viability. Sperm count decreases dramatically between 18 and 25 months in C57Bl/6 mice^24^ and the HCD carnosine is able to preserve sperm morphology in an accelerated aging senescent mouse model^25^. However, N-acetylcarnosine did not affect sperm count, or the percentage of live sperm in 24-month-old mice (Supp. Figure 4A).

Carnosine and other HCDs have also been implicated in improving neurologic function in aged animal and human models^26–28^. Thus, we performed a wide variety of assessments to examine the influence of N-acetylcarnosine treatment on the neurological system. N-acetylcarnosine treatment did not have any effects on brain size in either sex (Figure 3E, Supp. Figure 4B). The behavioral assessments also did not show differences in memory (fear conditioning and Y-maze) (Figure 3F&G, Supp. Figure 4C,D,F) or pain perception (mechanical or heat)^29^ (Figure 3H, Supp. Figure 4E&G) in male or female mice.

As one of the organ systems critical for survival we next profiled the influence of N-acetylcarnosine on cardio-renal phenotypes. N-acetylcarnosine treatment did not alter relative heart weight (Figure 4A, Supp. Figure 5A) or histology (Figure 4B, Supp. Figure 5H). There was a modest trend for lower heart rates in female mice (Figure 4C), suggesting a potentially positive effect of N-acetylcarnosine on heart or perhaps vagal tone. Fractional shortening (FS) in female N-acetylcarnosine-treated hearts was lower compared to the vehicle-treated mice (Figure 4F) with no changes to cardiac output or ejection fraction (Figure 4D&E). The lower FS was driven by ∼17% greater systolic left ventricular internal diameter compared to vehicle mice (p=0.002, data not shown), which suggests the N-acetylcarnosine treated mice are able to achieve similar blood delivery as the vehicle-treated mice with less forceful systolic contraction. This could indicate that the hearts from N-acetylcarnosine treated female mice have improved ventricular filling or are pumping blood against a lower afterload (systemic blood pressure). Metrics of diastolic function (E/A – indicative of filling velocity across the mitral anulus and E/e’ – indicative of late-diastolic ventricular filling) were not different in female mice (Figure 4G&H). Acetylcholine-evoked, endothelium-dependent vasodilation assessed using isometric myography was similar in vehicle and N-acetylcarnosine-treated mice (Figure 4I). However, addition of the endothelial nitric oxide synthase (eNOS) inhibitor L-NMMA attenuated vasorelaxation to a greater extent in vessels from N-acetylcarnosine compared to vehicle-treated mice (Figure 4I), a result that was substantiated by arteries from N-acetylcarnosine mice displaying a greater calculated nitric oxide bioavailability compared to vehicle mice (Figure 4J). Vascular smooth muscle responses were similar between groups (not shown). These findings are consistent with previous work that HCDs promote endothelium-dependent vasodilation^32,33^, and might have implications concerning systemic blood pressure.

**Figure 4.**
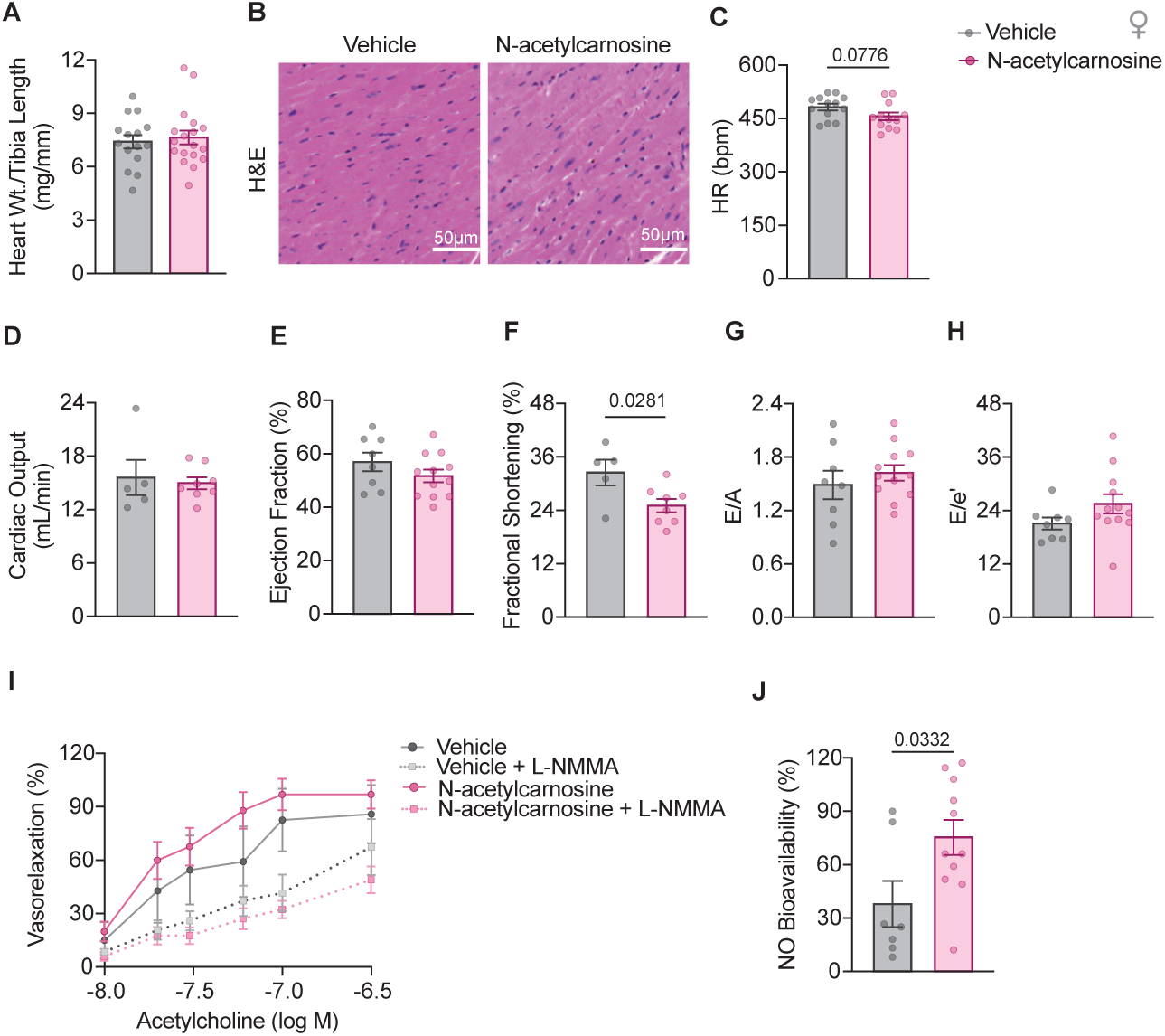
N-acetylcarnosine supplementation improves cardiovascular function in female mice. **A)** Heart weight normalized to tibial length was not different between groups of female mice. **B)** H&E staining of heart tissues was also indiscernible between groups. **C)** Heart rate measured via electrocardiogram was performed on anesthetized mice during echocardiography and was not different between groups. Metrics of systolic function including cardiac output **(D)** and ejection fraction **(E)** were not different between groups whereas fractional shortening was reduced by N-acetylcarnosine treatment **(F)**. Measures of diastolic function including E/A (ventricular filling) **(G)** and E/e’ (marker of left ventricular filling pressures) **(H)** were not different between groups. Isolated femoral arteries (n=8 vehicle, n=15 N-acetylcarnosine, 2 segments per mouse) were pre-contracted to 65% of maximal developed tension, and vasorelaxation to cumulative doses of Ach was recorded as percent (%) vasorelaxation. No differences were observed between groups. A second Ach dose-response curve was completed (30-min later) in arteries from both groups that incubated with LNMMA for 30-min. While L-NMMA attenuated vasorelaxation in arteries from vehicle and N-acetylcarnosine-treated mice, the contribution from NO to vasorelaxation was greater in the latter group **(I)**, an observation that was substantiated by calculating NO bioavailability **(J)**. Data are represented as Mean ± SEM. Statistical significance was assessed via two-way repeated measures ANOVA with Bonferroni correction, or unpaired t-test where appropriate. Comparisons were considered statistically significant if *p<*0.05. Heart Rate; HR.

N-acetylcarnosine appeared to influence heart to a lesser extent in male mice (Supp. Figure 5B-F), except for improved E/e’ indicative of late-diastolic ventricular filling (Supp. Figure 5G). It is important to point out that the N-acetylcarnosine treatment in our study started when mice were 18-months old, an age when some of the age-related decline in heart function had already began^30^. Thus, it remains possible that the treatment may have had a more substantial effect if initiated earlier. This is particularly the case as HCDs have been shown to improve excitation-contraction coupling via regulating calcium handling^31^. Overall, N-acetylcarnosine supplementation may improve some aspects of systolic function and systemic blood pressure regulation in aging female mice and diastolic function in aging male mice.

The cardiovascular system intricately interacts with the kidney to control blood pressure and filtration, a function that is known to decline drastically with aging. HCDs have been implicated to be protective of the kidney in animal models of obesity^18^. Nonetheless, N-acetylcarnosine treatment did not appear to influence kidney weight (Figure 5A), histology (Figure 5B), urine output (Figure 5C), creatinine clearance (Figure 5D), or glomerular filtration rates (GFR) (Figure 5E) in either sex (Supp. Figure 6A-G). These findings are particularly interesting given that other models of HCD enrichment such as whole-body deletions of carnosinase 1 (CN1) demonstrate preferential HCD enrichment in the kidney^34^. However, the GFR for both vehicle and N-acetylcarnosine treated mice are similar to GFRs that of healthy young mice we reported recently^35^ suggesting that kidney dysfunction may not have manifested by 24 months of age.

**Figure 5.**
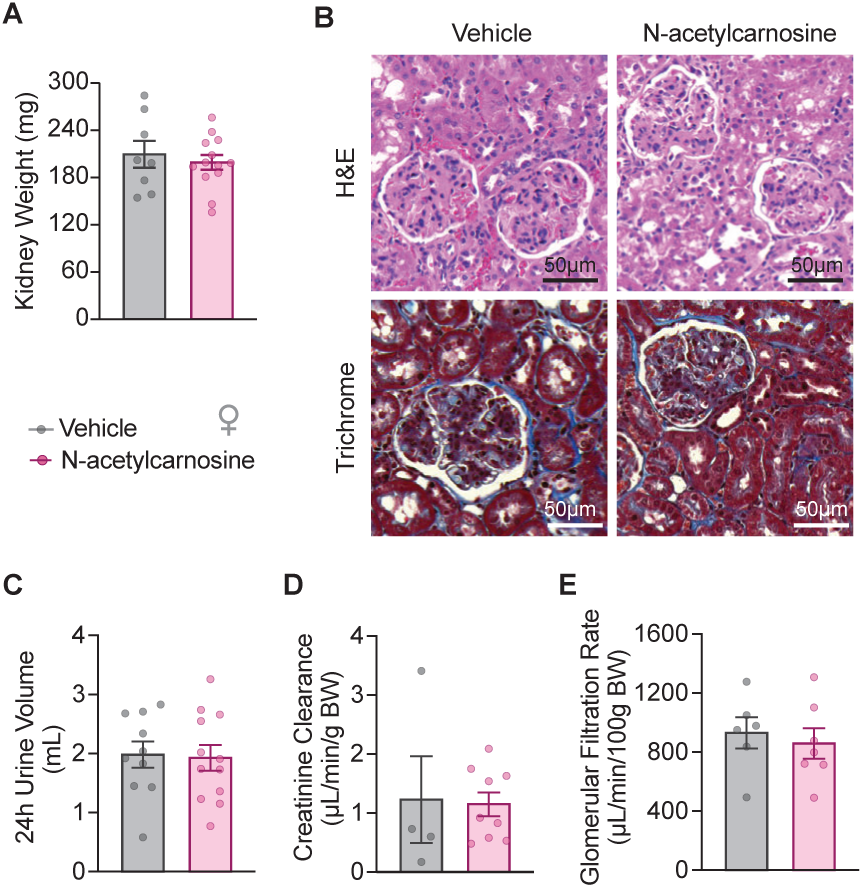
N-acetylcarnosine supplementation does not alter renal function in female mice. Kidney weight was not different between groups **(A)**. H&E and Trichrome stains of kidney sections were unremarkable and indistinguishable between groups **(B)**. Mice were placed in single housed, urine collection chambers for 24 hours to collect 24-hour urine volume which was not different between groups **(C)**. 24-hour urine and plasma and urine creatinine concentration were used to determine creatinine clearance **(D)** which also was not affected by N-acetylcarnosine. Glomerular filtration determined by monitoring the filtration of a FIT-c labeled senestrin under anesthesia was also not different between groups in female mice (n=5 vehicle, n=7 N-acetyl carnosine) **(E)**. Data are represented as Mean ± SEM. Statistical significance was assessed via two-way repeated measures ANOVA with Bonferroni correction, or unpaired t-test where appropriate. Comparisons were considered statistically significant if *p<*0.05.

Frailty is a hallmark of aging that very closely predicts mortality. Physical function, supported by the musculoskeletal system, is known to drastically become impaired in the process of aging to contribute to the decline in quality of life. Previously, we showed that N-acetylcarnosine robustly attenuates muscle atrophy and weakness induced by 1-week of disuse in older mice^14^. However, sarcopenia occurs gradually over a longer period, so the current study provided a perfect opportunity to determine the effects of N-acetylcarnosine treatment on loss of physical function with aging. N-acetylcarnosine did not appear to strongly influence muscle mass (Figure 6A) or myofiber cross-sectional area in fast-twitch extensor digitorum longus (EDL) (Figure 6B&C) or slow-twitch soleus muscles in either sex (Supp. Figure 7A-F). It is known that while C57Bl/6 mice begin to experience atrophy at 24 months, loss of muscle mass at peaks at about 30 months of age^36^ so this perhaps was not too surprising. In contrast, muscle weakness precedes muscle atrophy in sarcopenia^36^. Indeed, ex vivo assessment of skeletal muscle force-generating capacity revealed consistent increase in both EDL and soleus muscle strength induced by N-acetylcarnosine treatment in female mice (Figure 6D&E), but not in male mice (Supp. Figure 7G&H). A robust enrichment of N-acetylcarnosine in skeletal muscle in female mice, but not male mice (Figure 1C) is consistent with these observations. Nonetheless, these improvements were insufficient to be reflected in the *in vivo* assessments of physical function (Figure 6F-H, Supp. Figure 7I-K). These data implicate N-acetylcarnosine as a potentially viable strategy to delay features of sarcopenia in mice.

**Figure 6.**
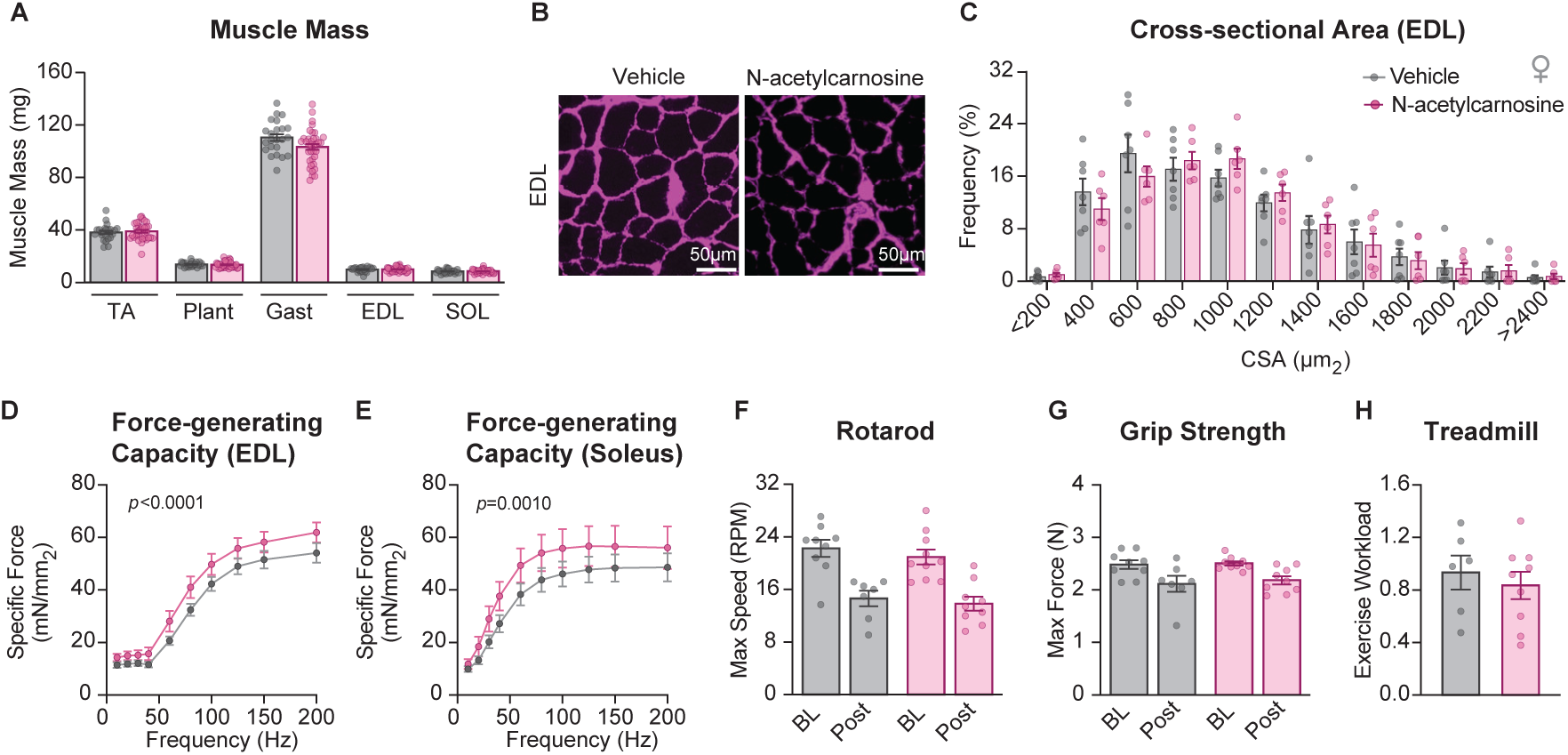
N-acetylcarnosine supplementation improves muscle function in female mice. **A)** Lower hind limb weights were not different between groups (n=23-24 vehicle, n=35-37 N-acetylcarnosine) with the exception of gastrocnemius (Gast). **C)** Histogram of the frequency of fibers sizes was not different between groups in EDL (n=6 vehicle, n=8 N-acetylcarnosine). **D)** Representative image of immunofluorescent stain for laminin used to determine muscle cross sectional area in the EDL. E*x vivo specific* force measured in EDL **(E)** (n=8 vehicle, n=8 N-acetylcarnosine) and SOL **(F)** (n=8 vehicle, n=9 N-acetylcarnosine) were elevated in N-acetylcarnosine-treated mice across stimulation frequencies ranging from 10-200 Hz. *In vivo* functional data as quantified by rotarod **(G),** grip strength **(H)**, and treadmill work capacity **(I)** were unaffected by N-acetylcarnosine treatment. Data are represented as Mean ± SEM. Statistical significance was assessed via two-way repeated measures ANOVA or two-way ANOVA with Bonferroni correction, or unpaired t-test where appropriate. Comparisons were considered statistically significant if *p<*0.05. Tibialis anterior; TA, plantaris; Plant, gastrocnemius; Gast, extensor digitorum longus; EDL, soleus; SOL.

Aging also profoundly influences physical function by weakening bones, particularly in post- menopausal women that experience osteoporosis. Micro-CT of femurs (Figure 7A) revealed that N-acetylcarnosine treatment increased trabecular number (Figure 7B, Supp. Figure 8A) but decreased trabecular spacing (Figure 7C, Supp. Figure 8B). This in parallel with unaltered trabecular thickness (Figure 7D, Supp. Figure 8C) led to increased bone volume fraction (Figure 7E, Supp. Figure 8D) in both sexes in the distal femur. In contrast, neither tibial trabecular bone nor inflammation at the knee joint were affected by N-acetylcarnosine treatment (Supp. Figure 8E-G). These data indicate an ability of N-acetylcarnosine to enhance bone density in the femurs of aging mice.

**Figure 7.**
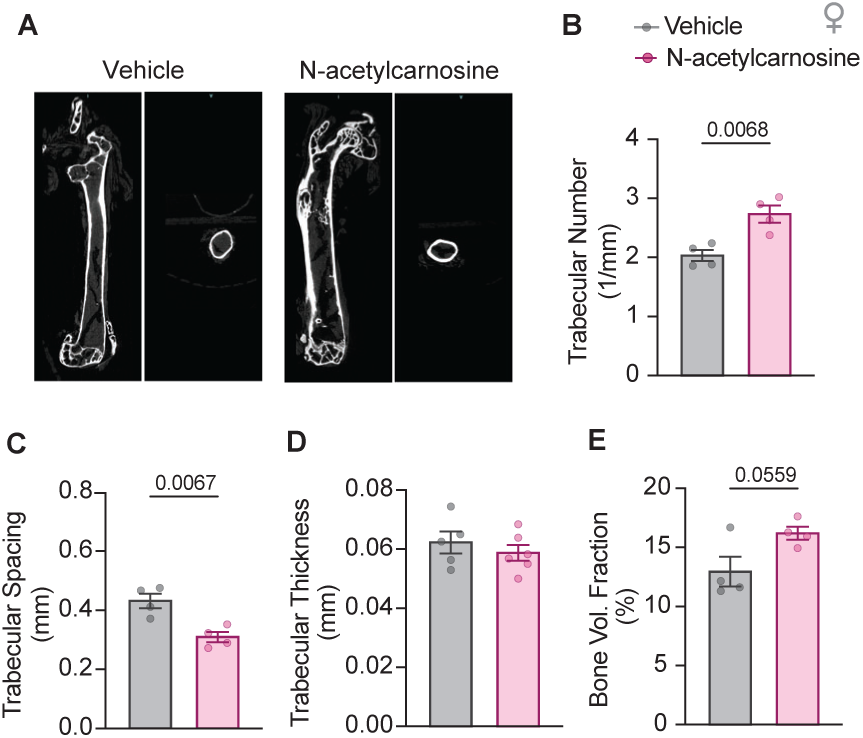
N-acetylcarnosine supplementation improves bone density in female mice. **A)** Representative µCT image of femur analyzed for bone density in the distal femur. µCT of femurs revealed increased trabecular number **(B)** and decreased trabecular spacing **(C)** without a change in trabecular thickness **(D)** which contributed to a near-significant increase in the proportion of bone volume **(E)** in the ROI. Data are represented as Mean ± SEM. Statistical significance was assessed via unpaired t-test where appropriate. Comparisons were considered statistically significant if *p<*0.05. Cross sectional area; CSA, micro computed tomography; µCT.

## Discussion

Aging is associated with exacerbated oxidative stress, however strategies broadly targeting reactive oxidative species (ROS) have been disappointing. This may be related to the presence of many primary and secondary ROS-related species and the important roles some of these species play in physiological signaling and hormetic response to stress. Carbonyls may be a more promising target given they are secondary ROS species, their formation is proportional to their metabolite precursors which may be altered in a context-dependent manner, and they are canonically not known to possess physiological signaling roles. Supplementation with HCDs such as N-acetylcarnosine, carnosine, their precursor β-alanine, and synthetic analogs have established efficacy for mitigating age-related pathologies such as sarcopenia^14,15^, insulin resistance^12,21,22^, cardiovascular^33,37,38^ and renal dysfunction^39^ primarily in contexts outside of aging purportedly via scavenging of reactive carbonyls. Ours is the first study to simultaneously investigate the effects of N-acetylcarnosine on these systems during aging *per se* in a longitudinal study.

Supplementation with N-acetylcarnosine improved survival of female mice by approximately 50%. While the mechanism by which female mice preferentially benefited is unclear, certain tissues from female N-acetylcarnosine-treated mice that displayed sex-specific effects, were more enriched for N-acetylcarnosine (Figure 1C). For example, N-acetylcarnosine was only significantly elevated in gWAT from female N-acetylcarnosine-treated mice (Supp. Figure 1). In parallel, we observed a disparate effect of N-acetylcarnosine to promote weight retention in female mice which appeared to be driven by larger gWAT (Figure 2G) without compromising metabolic function. A finding in line with previous work demonstrating the ability of HCDs to protect against obesity-related glucose intolerance^12,13,19^. While, the mechanism of this adipose retention is unclear, our indirect calorimetry and activity data demonstrate decreased energy expenditure and activity in female N-acetylcarnosine treated mice. Interestingly, recent research suggests that decreased metabolic rate induced by a torpor-like state increases survivability and delays aging in mice^40^. Carnosine treatment has also been demonstrated to mitigate adipocyte senescence via scavenging of 4-HNE in aging mice^41^ which could promote retention of fat mass as we observed in the N-acetylcarnosine-treated mice.

The carbonyl scavenging ability of lipid carbonyls such as 4-HNE may have also contributed to vascular, muscle, and bone function as well. 4-HNE uncouples NO production in endothelial cells^42^ leading to impaired NO bioavailability and carnosine has been shown to facilitate NO production *in vitro*^33^. While this increased NO bioavailability we observed in the N-acetylcarnosine treated mice did not enhance endothelium-dependent vasodilation *ex vivo*, NO bioavailability is closely associated with vascular health and appropriate blood pressure regulation which we did not measure. Improved NO bioavailability may also protect old mice from additional insults (e.g. ischemia reperfusion, dyslipidemia etc.) which should be tested in future experiments. It is worth noting that N-acetylcarnosine drastically reduced protein carbonyl burden in the heart, even though N-acetylcarnosine treated-mice did not exhibit overt improvement in heart function discerned from echocardiography. However, heart functional decline likely occurs earlier than 18 months which may have limited the therapeutic capacity of N-acetylcarnosine in this study.

We previously found N-acetylcarnosine was effective at mitigating 4-HNE modified proteins in muscle and preserved muscle force following 7-days of hindlimb unloading. Here, we demonstrate that N-acetylcarnosine was effective at preserving *ex vivo* muscle force production during aging.

HCDs have been shown to chelate and alter the concentration of di-cations such as calcium which may be an alternate mechanism to carbonyl sequestering that may have contributed to this observation by altering cross bridge kinetics. This effect did not result in improved functional metrics (i.e. treadmill, grip strength, rotarod) but these measurements mostly rely on submaximal strength, do not necessarily reflect functional capacity relevant for trip and fall prevention (important health factors for elderly individuals). Bone forming osteoblasts cells are also negatively impacted by oxidative stress^43^ which may have been attenuated by N-acetylcarnosine treatment to increase femur trabecular bone density. Improved bone density is also translationally relevant for older individuals with increased risk of bone fractures which can dramatically increase morbidity and mortality especially in women. While the collection of these physiologic improvements does not provide a clear explanation of why the female N-acetylcarnosine mice improved survival by 50%, we speculate that the retention of adiposity may have broadly delayed frailty onset which manifested with improved NO bioavailability, muscle force capacity, and bone density.

In conclusion, N-acetylcarnosine treatment improves functions of multiple organ systems to increase survival in female mice. As an endogenously present metabolite, N-acetylcarnosine likely represents a safe, commercially scalable, and economically affordable treatment in humans. The current intervention spanned 18-24 months of age, and it remains possible that N-acetylcarnosine treatment has additional protective effects on age-related declines that occur outside this window. Humans are known to have substantially different HCD metabolism and pharmacokinetics than mice. Thus, N-acetylcarnosine is a strong candidate to be considered for clinical testing to determine its efficacy in mitigating age-related health decline in humans.

## Methods

### Mice

All experiments were performed at the University of Utah and were approved by the Institutional Animal Care and Use Committee. C57Bl/6 mice were ordered from the National Institutes on Aging’s aged mouse colony at 17-months of age and fed a normal chow diet, and housed with a 12-hour light/dark cycle in a temperature-controlled room for the duration of the study. Mice were ordered in cohorts of 20 mice per months with three cohorts consisting of 10 males and 10 females, and five subsequent cohorts consisting of 20 female mice after observing certain trends in female mice only. Upon arrival mice were weighed, counterbalanced by weight, and re-caged to five mice per cage when possible and needed. Mice were monitored for the next three weeks with minimum contact to allow them to acclimate to their new environment. After three weeks of acclimation, baseline measures of body composition, grip strength, rotarod, and Y-Maze were taken (described below).

### N-acetylcarnosine Treatment

Cages of mice were randomized to receive normal drinking water (vehicle) or drinking water supplemented with 80 mM N-acetylcarnosine prepared as previously described^14^. Briefly, tap water was autoclaved and supplemented with N-acetylcarnosine to final concentration of 80 mM. Both vehicle and N-acetylcarnosine water were titrated to a final pH of 7.5. Water was prepared fresh and replaced weekly. At the time water is replaced, the mice are weighed, fresh bedding is provided, and food and water consumption are measured. The intervention begins 4 weeks after the mice have been acquired at 80 weeks of age and continued for 24 weeks. Over the course of the intervention, mice are also monitored for general well-being and treated as needed by the veterinarians. If deemed necessary by the veterinarians, mice were euthanized prior to the end point. Reasons for early euthanasia included unresolving scruff wounds, anal prolapse, enlarged tumors, and extreme weight loss. In addition to weekly routine monitoring, mice were assessed every 8 weeks for body composition. Four weeks prior to the study endpoint, physiological endpoint measures (body composition, grip strength, rotarod, treadmill test, Y-maze, Comprehensive Laboratory Animal Monitoring System (CLAMS), echocardiography, 24-hour urine collection, glomerular filtration rate, fear conditioning, Von Frey test, hot plate test, and intraperitoneal glucose tolerance test). This timing was chosen to limit the proximity of tests to one another to limit stressing the mice, and to prevent any resultant stress from confounding terminal tissue collection. To limit measurement bias, the tester performing these assessments was blinded and in the case of repeated measures, the same tester performed the assessment at all time points.

### Body Composition

Body composition was assessed at the beginning of the intervention and 8, 16, and 24 weeks of treatment via Bruker Minispec NMR (Bruker, Germany).

### Grip Strength

Whole body grip strength was assessed at baseline and between week 100 and 103 of the intervention utilizing a grip strength meter with a mesh wire attachment (Columbus Instruments, Columbus, OH, USA) with methods modified from^44^. Briefly mice were acclimated to the testing room for 30 minutes. The tester then removed a mouse from the cage (blind from the group allocation of that cage) and placed it onto the mesh wire attached to the dynamometer. Gripping at the base of the tail, the mouse was pulled steadily, with its torso horizontal to the ground, back towards the tester. This trial was repeated three times with 30 seconds between each bout and the highest force being reported.

### Rotarod

Rotarod assessment was performed at baseline and between week 100 and 103 of the intervention as previously described^44^. Briefly, mice were trained on the Rotamex-5 (Columbus Instruments, Columbus, OH) for three days prior to testing by placing mice on the apparatus at a constant speed of 4 rpm for 5 minutes, replacing them if they were to fall. On the day of testing, mice were acclimated to the testing room for 30 minutes prior to testing. After acclimating, mice were placed on the apparatus and the instrument was run at 4 rpm, accelerating 0.3 rpm/s. Mice were allowed to continue if they were able to grip the rotarod to prevent a fall. Timer continued until the mice fell to the bottom. Once the mice were assayed, the test was repeated starting with the first mouse assayed until all mice were assayed three times with the best score being reported.

### Treadmill Exercise Capacity

The treadmill test was performed between week 100 and 103 of the intervention as previously described^30^ prior to terminal experiments. Every day for the three days prior to the treadmill exercise test, mice were acclimated to the treadmill by placing each mouse in the treadmill for 5 minutes with the belt being stationary followed by 5 minutes with the belt running at 5 meters per minute. On the day of the test, mice were acclimated to the test room for 30 minutes. Four mice at a time were placed on the treadmill with walls separating adjacent mice. Mice began by running at 5 meters per minute at a 25% grade for 1 minute. The speed was increased by 1 meter per minute every minute until maximal exercise capacity was achieved. The mice were encouraged to run by tapping the mice with a wire cleaner brush. Maximal capacity was determined by a mouse’s inability to run on the treadmill following 5 consecutive coercions with the wire brush followed by 2 attempts to place the mouse in the middle of the treadmill by hand. Maximal exercise capacity was calculated as follows:

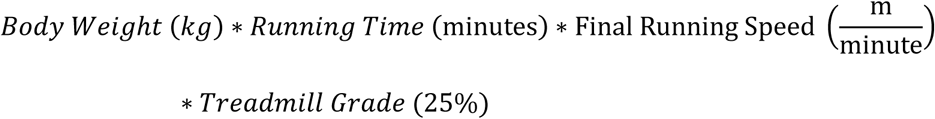

### Y-Maze

Y-Maze was performed at baseline and between week 100 and 103 of the intervention. Mice were acclimated to the testing room 30 minutes prior to the assessment. The Y-maze was placed on the floor in the same orientation in the same room for repeated measures. A camera was suspended above the Y-maze apparatus to record the test. The mouse was then placed in the center for 20 seconds of the maze with three walls blocking the path to the maze arms to acclimate to the apparatus. After acclimating to the apparatus, video recording was initiated and the blockades to the maze arms were removed. The branches that the mouse traveled to was recorded for 15 branches unless the mouse returned to the same arm twice in a row at which point the test was terminated. Spontaneous alternations were calculated by counting the number of unique groupings of three arms visited across the entire test.

### CLAMS

Comprehensive laboratory animal metabolic system (CLAMS, Columbus Instruments, Columbus, OH) was performed between week 100 and 103 of the study to measure energy expenditure (VO_2_), respiratory exchange ratio (RER; VCO_2_/ VO_2_), food and water consumption, and physical activity.

### Echocardiography

Echocardiography was performed between week 100 and 103 of the study. Mice were induced and maintained under light anesthesia by inhalation of isofluorane. Cardiac function was measured using a VisualSonics Vevo 2100 ultrasound machine with MS400 probe. B- and M- mode sequences were acquired in the parasternal long- and short-axis orientations and used to assess systolic function and to quantify morphological features. Mitral inflow velocities (E and A), and early diastolic mitral annulus tissue velocity (e’) were measured in the apical, four-chamber view using flow and tissue doppler respectively. These values were used to assess diastolic function based on E/A and E/e’ ratios. Echocardiographic data was analyzed using VisualSonics VevoLab.

### 24-Hour Urine Collection and Creatinine Clearance

A 24-hour urine collection for determination of creatinine clearance was performed between week 100 and 103 of the study. Mice were placed in an MMC100 Metabolic Cage for 24 h at which point urine and cheek blood were collected. Body weight, urine volume, food, and water intake were all observed throughout the experiment. Mice were fed with a standard chow gel diet with free access to water. Urine samples were centrifuged at 21,000 x g for 15 min, then aliquoted, and stored at -80 C. Urine and plasma creatinine were measured using QuantiChrom Creatinine Assay Kit, BioAssay Systems, Hayward, CA (Cat. No. DIUR-500) according to the manufacturer’s instructions. Creatinine clearance was calculated as follows:

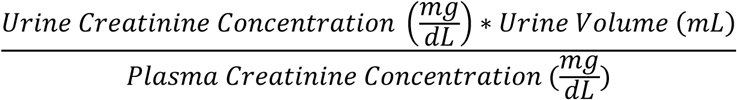

### Glomerular Filtration Rate

Glomerular filtration rate (GFR) was measured between week 100 and 103 of the study as previously described^45^. Mice were injected with fluorescein isothiocyanate-senestrin (Medibeacon, Mannheim, Germany) retro-orbitally (7.5 mg/100 g body weight). The NIC-Kidney (Medibeacon) was used to detect fluorescence in the skin on the shaved back over 1 h. GFR was calculated based on the kinetics of fluorescence decay as previously reported^46^.

### Fear Conditioning

Fear conditioning was conducted between week 100 and 103 of the study using a med-associates sound-attenuated chamber (29x25x26 cm) with no scent, plexiglass roof and front door, aluminum side walls, a plastic black wall and fitted with a top-down camera (*Context A)* (Med Associates; St. Albans, Vermont). Conditional and contextual memory were assessed on two subsequent days. On day 1 mice were acclimated to the test room for at least 1 hour. After shock calibration, an individual mouse was placed in the chamber for 3 minutes. At 120 seconds, a tone was delivered for 20 seconds. Two seconds prior to termination of the tone, a 0.3 mA foot shock was delivered lasting 2 seconds. The mouse remains in the chamber until the end of the 3 minutes. The mouse is then returned to its cage, and the chamber is cleaned with 70% ethanol before testing the next mouse. On day 2, contextual fear conditioning is tested by introducing mice to *Context A* for 3 minutes without tone or shock. The percent time freezing on day 1 and day 2 are reported as metrics of contextual memory. One hour later, mice are tested for conditional memory by being introduced to a new context which differs from *Context A* by placing curved plastic board inside the chamber, a plastic board underneath the grid floor, and the chamber is scented with vanilla extract (*Context B)*. Mice are allowed to explore for 3 minutes without foot shocks but with the same tone as on the first day occurring during the last minute. The percent change (%Δ) of time freezing before and after the tone on day 1 compared to day 2 is reported as the metric for conditional memory.

### Von Frey Test

The Von Frey test was performed between 100 and 103 weeks of the study. Mice were acclimated to the procedure room for 30 minutes prior to testing. Mice were placed in a small chamber with grid wire flooring. The Von Frey apparatus was fitted with three filaments of different tensions. These filaments were applied to the plantar region of the mouse’s foot and the force needed to elicit withdraw of the foot is recorded. Each test was repeated three times with 15 minutes between each test.

### Hot Plate Test

The hot plate test was performed between 100 and 103 weeks of the study. Mice were acclimated to the procedure room for 30 minutes prior to testing. A glass enclosure was warmed to 50 degrees Celsius and a camera was affixed to record the animal’s behavior in response to being placed in the enclosure. Change in behavior indicating pain (e.g. picking up paws, rearing etc.) was scored and timed by 3 independent and blinded individuals.

### Terminal Biological Sample Collection

On week 104, 4-hour fasted mice were anesthetized via intraperitoneal injection of 80 mg/kg ketamine and 10 mg/kg xylazine. Hindlimb muscles (Tibialis anterior [TA], Extensor digitorus longus [EDL], soleus, plantaris, and gastrocnemius) were quickly collected from live mice, weighed, and either tied with sutures on distal and proximal tendons for *ex vivo force* (EDL and soleus; described below), flash frozen in liquid nitrogen, or embedded in OCT and frozen in liquid nitrogen-cooled isopentane for histological assessment (described below). After collecting muscles, blood was collected via cardiac puncture followed by collecting, weighing, preservation of the remainder of the tissues (heart, liver, kidney, inguinal white adipose tissue [iWAT], gonadal [gWAT], brain, sperm, and bones).

### Targeted Metabolomics

Plasma, urine, muscle (TA), whole brain, heart, iWAT, gWAT, kidney, and liver were processed from frozen and assessed for N-acetylcarnosine, carnosine, histidine, anserine, and homocarnosine concentrations. Sample preparation and metabolomics methods were based on recently published protocols^4^.

Frozen tissues were quickly weighed and immediately placed in HPLC-grade water with 10 mM HCl (12 μL extraction solution per mg tissue) followed by homogenization in a bead mill homogenizer at 4°C. Homogenates were centrifuged (20 min, 3000 × g, 4°C) and the resulting supernatants were immediately diluted in a 3:1 ratio with ice-cold acetonitrile. The mixture was vortexed for 15 s and kept on ice for 15 min. After a second centrifugation step (12 min, 20,000 × g, 4°C), samples were stored at −80°C until the day of analysis. Plasma and urine samples (37.5% of total mixture) were combined with acetonitrile containing 1% formic acid (54%) and HPLC-grade water (8.5%). Samples were vortexed for 15 s and centrifuged (15 min, 15,000 × g, 4°C), after which supernatant was collected and stored at −80°C until the day of analysis.

Targeted metabolite measurements were performed on a triple quadrupole LC-MS instrument (Agilent 6470). Normal phase chromatography was used to separate polar metabolites using a Luna 5µm NH_2_100 Å LC column (Phenomenex 00G-4378-E0) equipped with a SecurityGuard cartridge (KJ0-4282, Phenomenex). Injection volume was 7 µL, except for urine (4 µL). Mobile phases consisted of Buffer A (water with 10 mM ammonium formate) and Buffer B (95:5 acetonitrile:water with 10 mM ammonium formate). Each run started 100% B for 2 minutes, followed by a gradient starting at 100% B changing linearly to 50% A and 50% B over the course of 10 minutes, followed by 50% A and 50% B for 3 minutes, and re-equilibration at 100% B for 2 min (flow rate 0.7 mL/miN). MS analysis was performed using AJS in positive mode with the following parameters: dry gas temperature 250 °C, gas flow 12 L/min, nebulizer pressure 25 psi, sheath gas temperature 300 °C, the sheath gas flow 12 L/min, capillary voltage 3,500 V. Multiple reaction monitoring was performed to detect N-acetylcarnosine (269.1 → 110.1 *m/z*), carnosine (227.1 → 110.1 *m/z*), anserine (241.1 → 109.1 *m/z*), homocarnosine (241.1 → 110.1 *m/z)*, histidine (156.1 → 110.1 *m/z*). Secondary transitions^4^ were used as confirmation of the peak identity. Fragmentor voltage for the various metabolites was 80-100, collision energy 10-24 V. Endogenous metabolite concentrations were quantified using an external standard curve with known concentrations analyzed alongside the biological samples in the same run, except for homocarnosine. Total peak areas were used to calculate metabolite concentrations within each sample.

### Protein Carbonyl Assay

Protein carbonyls were measured via DNPH derivatization of proteins from tissues lysed in 50 mM MES buffer pH 7.4, 50 mM EDTA via protein carbonyl colorimetric assay kit (10005020, Caymen Chemical, Ann Arbor, MI).

### *Ex Vivo* Muscle Force

Force generating capacity of both the soleus and EDL were measured as previously described^14,47–49^. Briefly, soleus and EDL were tied with suture at proximal and distal tendons then attached to an anchor and force transducer in a tissue bath (Aurora Scientific, Model 801C) while being submerged in oxygenated Krebs-Henseleit Buffer solution at 37° C. The optimal length of the muscle was determined via maximum twitch force production. The buffer was then replaced with a fresh buffer solution and equilibrated for 5 minutes. After equilibration, a force-frequency sequence stimulated the muscle at increasing frequencies (10, 20, 30, 40, 60, 80, 100, 125, 150, and 200 Hz) with a 2-minute rest interval between each frequency. The rates of contraction and relaxation were quantified as previously described^14,47,48^. The Aurora Scientific DMAv5.321 software was utilized for analysis of force production data.

### Muscle Histology and CSA

OCT embedded soleus and EDL samples were mounted on a precooled platform and sectioned at 10 μm thickness with a cryostat (Microtome Plus). 6-8 serial sections from each sample were transferred onto the same slide. Sections then underwent blocking for 1 hour at room temperature with M.O.M. mouse IgG blocking reagent (Vector Laboratories, MKB-2213), after which they were incubated overnight at 4° C with a concentrated primary antibody targeting laminin (1:200, Sigma, L9393) in 2.5% horse serum in PBS. The next day, sections were incubated for 1 hour at room temperature with anti-rabbit IgG conjugated to Alexa Fluor 647 (1:250, Invitrogen, A21245). Sections were then washed three times in PBS, fixed with methanol for 5 minutes at room temperature and then washed 2 more times in PBS. Finally, slides were mounted with mounting medium (Vector 136 Laboratories, H-1000). Slides were imaged on a Zeiss Axioscan.Z1 at 20X magnification. Masks for fibers were generated via Cellpose (V3.0)^50^ and then imported into ImageJ software where CSA was quantified.

### Tissue H&E and Trichrome Staining

Portions of tissues (e.g. liver, heart) or antiliteral tissues (e.g. kidney) that were not flash frozen in liquid nitrogen, were taken from each mouse and immediately submerged in 4% paraformaldehyde for 12 hours and then 70% ethanol for 48 hours. Tissues were sectioned were then embedded in paraffin, sectioned at 10-μm thickness, stained for Masson’s Trichrome to assess fibrosis or hematoxylin and eosin (H&E) to determine fat droplet accumulation and general morphological features. Samples were imaged on Axio Scan Z.1 (Zeiss).

### *Ex Vivo* Vascular Function

Vasocontractile responses to potassium chloride (KCl, 20–100 mM) and phenylephrine (PE, 10^−8^- 10^−5^ M; not shown), and vasorelaxation responses to acetylcholine (10^−8^-10^−5^ M] and sodium nitroprusside (10^−9^-10^−4^ M; not shown), were completed using isometric myography as we have described^51^. To determine the contribution to vasorelaxation from endothelial nitric oxide synthase (eNOS), a second Ach dose-response curve was completed (30-min later) in the presence of the eNOS inhibitor NG methyl L arginine acetate salt (L-NMMA; 1mM]. To calculate NO bioavailability for each group, maximal vasorelaxation to Ach in the presence of L-NMMA was subtracted from maximal vasorelaxation to Ach in the absence of L-NMMA, and all values were averaged, and compared between groups. For all responses, values from two femoral artery segments per mouse were averaged.

### Knee Joint Histology and Scoring

To evaluate the effects of N-acetylcarnosine on OA development in aged mice right hind limbs were collected and fixed in 10% formalin. Bone microstructure analysis was performed as stated below. Following bone microarchitecture analysis by micro-CT joints were decalcified, dehydrated, and embedded in paraffin. The joint was sectioned at a thickness of 8 μm. To determine OA severity, joint sections were stained with hematoxylin, fast green, and Safranin-O. Three independent, blind graders scored joint sections for OA using a modified Mankin scoring system (Glasson, 2007). Scores were averaged among graders for the whole joint.

### Distal Femur Micro-Computed Tomography

Femur bone structure and morphological changes were measured using a Siemens Inveon® Multi-Modality microPET/SPECT/CT machine using the following imaging parameters: an X-ray tube potential of 80 kVp, an X-ray tube current of 100 μA, an integration time of 2600 ms, and an isotropic voxel size of 10 μm. Image analysis was performed using Inveon® Research Workplace (IRW) 4.0. A 100-slice volume of interest (VOI) was selected for the cortical bone, centered about the mid-diaphysis. Cortical bone was delineated from the medullary cavity using a fixed threshold that remained constant across specimens. A second 100-slice VOI was selected for the trabecular bone, beginning just proximal to the distal epiphyseal plate and extending into the metaphysis. Irregular anatomical contours were drawn slice-by-slice throughout this VOI just within the endocortical boundary as described by Bouxsein and colleagues^52^. Trabecular bone was delineated from the medullary cavity using a fixed threshold that remained constant across specimens. IRW’s *Bone Morphometry* tool was used to compute bone volume fraction, trabecular spacing, trabecular thickness, and trabecular number. The *Annotation and Measurement* tool was used to estimate the cortical thickness from four measurements at the mid-diaphysis (corresponding to the major and minor axes of an ellipse approximating the cross-section of the femur).

### Proximal Tibia Micro Computed Tomography

Tibial bone structure and morphological changes were measured in the intact hind limbs of mice by micro-CT (PerkinElmer Quantum GX2 microCT) with a 5.5-μm isotropic voxel resolution. Subchondral/trabecular regions were segmented using CTAn automatic thresholding software. The tibial and femoral epiphyses were selected using the subchondral plate and growth plate as references. The tibial metaphysis was defined as the 300 µm region directly below the growth plate. Micro-CT limb images were analyzed for bone volume/total volume (BV/TV), trabecular bone number (Tb.N), trabecular bone thickness (Tb.TH), trabecular bone separation (Tb.SP)^53^.

### Statistics

Significance for all statistical comparisons was set at *p*<0.05. Data were tested for normal distribution via visual inspection of QQ plots for linearity. Student’s T test or Wilcoxon Signed Rank test were used for comparison of two groups depending on normality of data. Repeated measurements, two-way ANOVAs or if some repeated measures were missing, mixed effects two-way ANOVAs were utilized to test group by time interactions with Bonferroni Post Hoc analyses applied where appropriate. Individual sample values as well as estimates of central tendency (mean or media where indicated) are presented in all bar graphs. Standard error of the mean is reported as the variance metric. Z-scores for radar plots in Figure 1 were calculated by considering the distribution of all samples regardless of sex or group. Statistical analyses including two-way ANOVA, mean, and SEM for these data are presented in Supp. Figure 1.

## Data Availability

Raw data will be available with reasonable request to the corresponding author.

## Ethics Declaration

The authors declare no competing interests.

## Author Contributions

ERM conceived of and designed the study, conducted experiments, performed data analysis, and wrote and edited the manuscript. KF conceived of and designed the study, analyzed data, and wrote and edited the manuscript. MJD and AC conceived of and designed the study, analyzed data, and edited the manuscript. JLS, SW, JS, SM, IYG, TS, MRG, SB, NSH, JDS, and JZL conducted experiments, analyzed data, and edited the manuscript. NYM, GN, BW, DS, JW, MJ, GH, TK, JKL, BR, KTO, NJB, HL, LSN, YL, CFD, JEC, CMK, NR conducted experiments and analyzed data.

## Acknowledgements

This work was supported by NIH grants DK110858 (JEC); HL167866, DK128819 (SB); AR083904, AR071967 (CMK); DK133271 (NR); NS137105 (JDS); AG065993, CA272529, CA286584 (AC); DK130641, MH133313, DK136526, OD034455, DK124265 (JZL); AG074535, AG076075 (MJD); AG074535, DK127979, DK107397, GM144613 (KF); HL007576 (ERM); DK137475 (MRG); AHA predoctoral grant AHA1025910, University of Utah Graduate Research Fellowship (SM); and the grant-in-aid for Japan Society for Promotion of Science (JSPS) Fellows 24KJ2039 (TK). Figures were edited by Nikita Abraham. This work was also supported by the following University of Utah Core Facilities: Cell Imaging Core, Preclinical Imaging Core, and Metabolic Phenotyping Core.

---

**Supplemental Figure 1.**
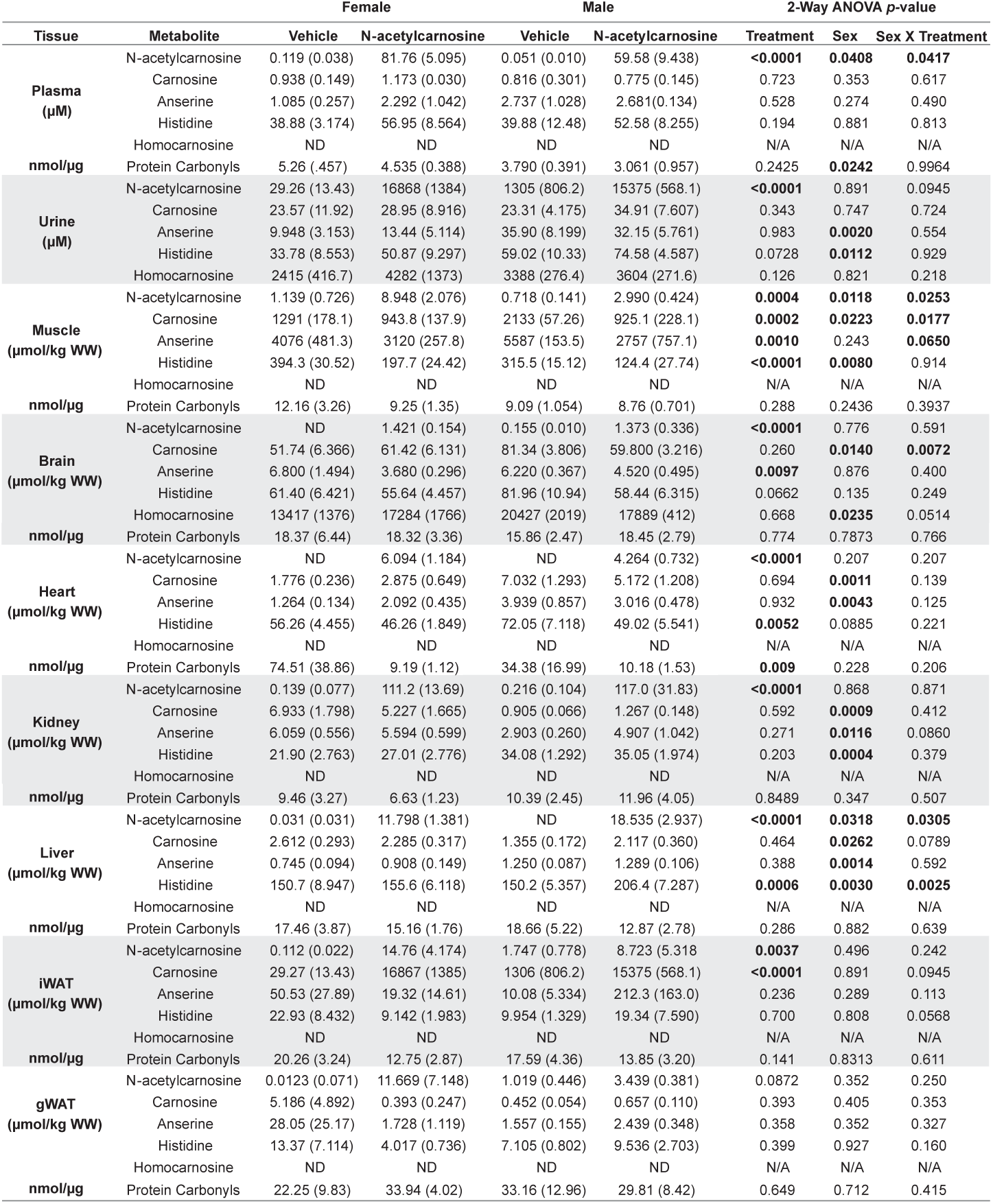
Histidine-containing dipeptides and protein carbonyl concentrations across tissues. Targeted metabolomics via LC-MS was utilized to quantify N-acetylcarnosine, carnosine, anserine, histidine, and homocarnosine in plasma, urine, muscle (TA – HCDs, Gastroc. – carbonyls), brain (cortex), heart (ventricle), kidney, liver, iWAT, and gWAT. Data are presented as Mean ± SEM. Two-way ANOVA was used to test statistical significance. The p-values for the main effects of N-acetylcarnosine treatment (Treatment), sex, and the interaction of these variables (sex x treatment) are reported in the last three columns respectively. Significance was set a *p*<0.05. Bolded *p-*values indicate a significant main effect.

**Supplemental Figure 2.**
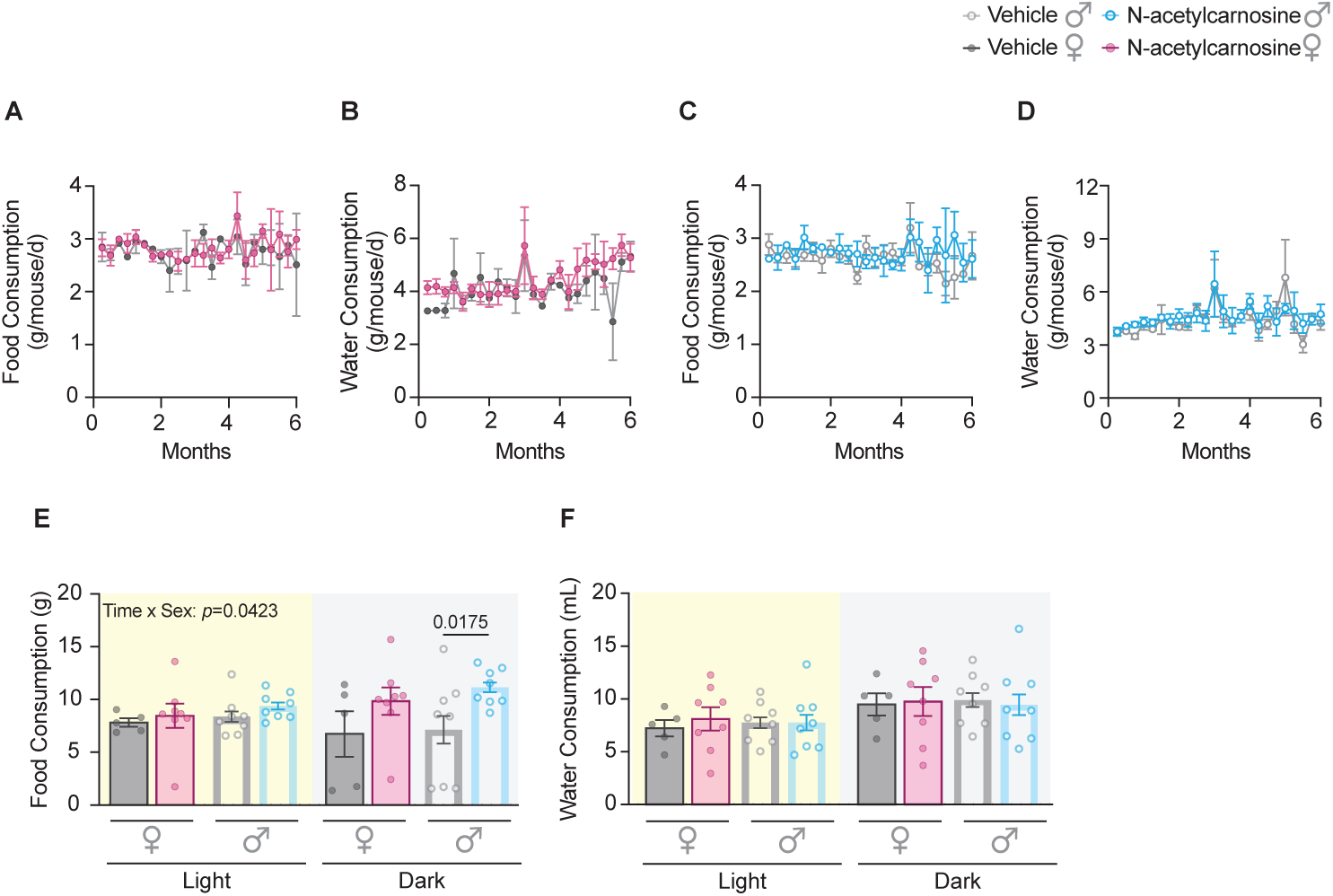
N-acetylcarnosine supplementation does not alter food or water intake. Free-living female food **(A)** and water **(B)** and male food **(C)** and water **(D)** were not different on a per cage basis. Data are represented as mean ± SEM of the grams of consumption per mouse per day. Food **(E)** and water **(F)** consumption was also determined during the CLAMS (n=8 vehicle, n=8 N-acetylcarnosine) assessment of single-housed mice. Food consumption was only significantly increased in male mice during the dark phase. Data are presented as Mean ± SEM. Statistical significance was assessed by two-way repeated measures ANOVA with Bonferroni post-hoc where appropriate. Significance was set to *p* < 0.05).

**Supplemental Figure 3.**
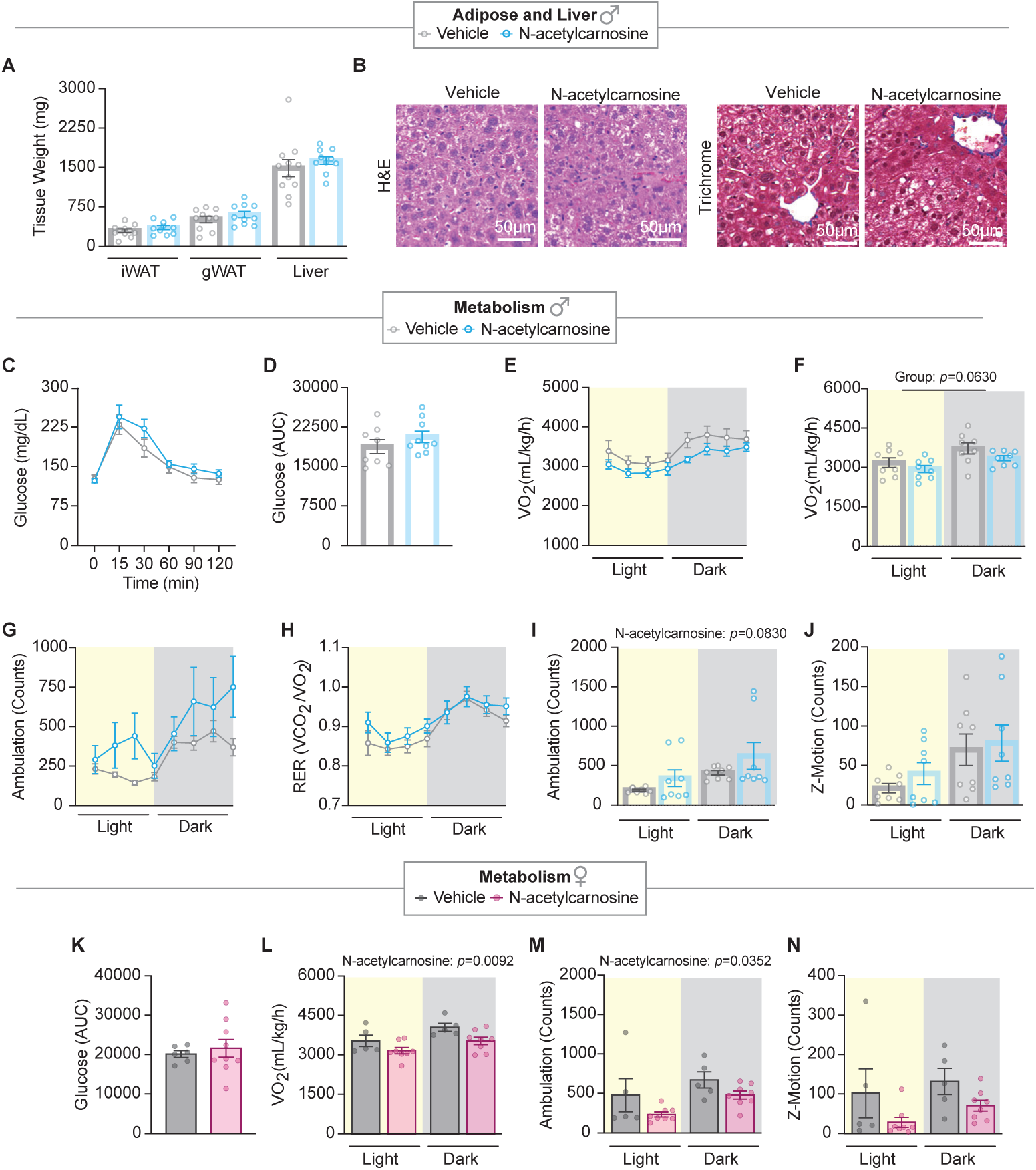
N-acetylcarnosine supplementation does not glucose tolerance and attenuates energy expenditure. **A)** Male iWAT, gWAT, and liver weights were not different between groups. **B)** H&E and Trichrome stains of liver sections in male mice were not remarkably different between groups demonstrating limited lipid infiltration and fibrosis in both groups. **C)** 2-hour IPGTT revealed no differences in plasma glucose in male mice at any time point or as represented as area under the curve (AUC) **(D)**. **E)** CLAMS revealed a trend for lower hourly energy expenditure (relative VO_2_), and a significantly lower energy expenditure AUC in the N-acetylcarnosine treated male mice in the light and dark time points **(F)**. **G)** Hourly ambulation (3h averages) and RER **(H)** in male mice was not different between groups during CLAMS. Ambulation AUC was significantly higher in the male, N-acetylcarnosine treated mice **(I)** in light and dark time points. **J)** Motion in the Z-direction was not affected by N-acetylcarnosine in the male mice. **K)** Female glucose AUC during 2-hour IPGTT was not different between groups. **L)** Female energy expenditure (relative VO_2_) AUC was lower in N-acetylcarnosine treated mice. **M)** Total ambulation was lower in light and dark time points in female N-acetylcarnosine treated mice. **N)** Z-motion frequency was not different between female vehicle and N-acetylcarnosine-treated mice. Data are presented as Mean ± SEM. Two-way RMANOVA or T-tests were used for statistical comparison with Bonferroni post-hoc test where appropriate. Comparisons were considered statistically significant if *p*<0.05.

**Supplemental Figure 4.**
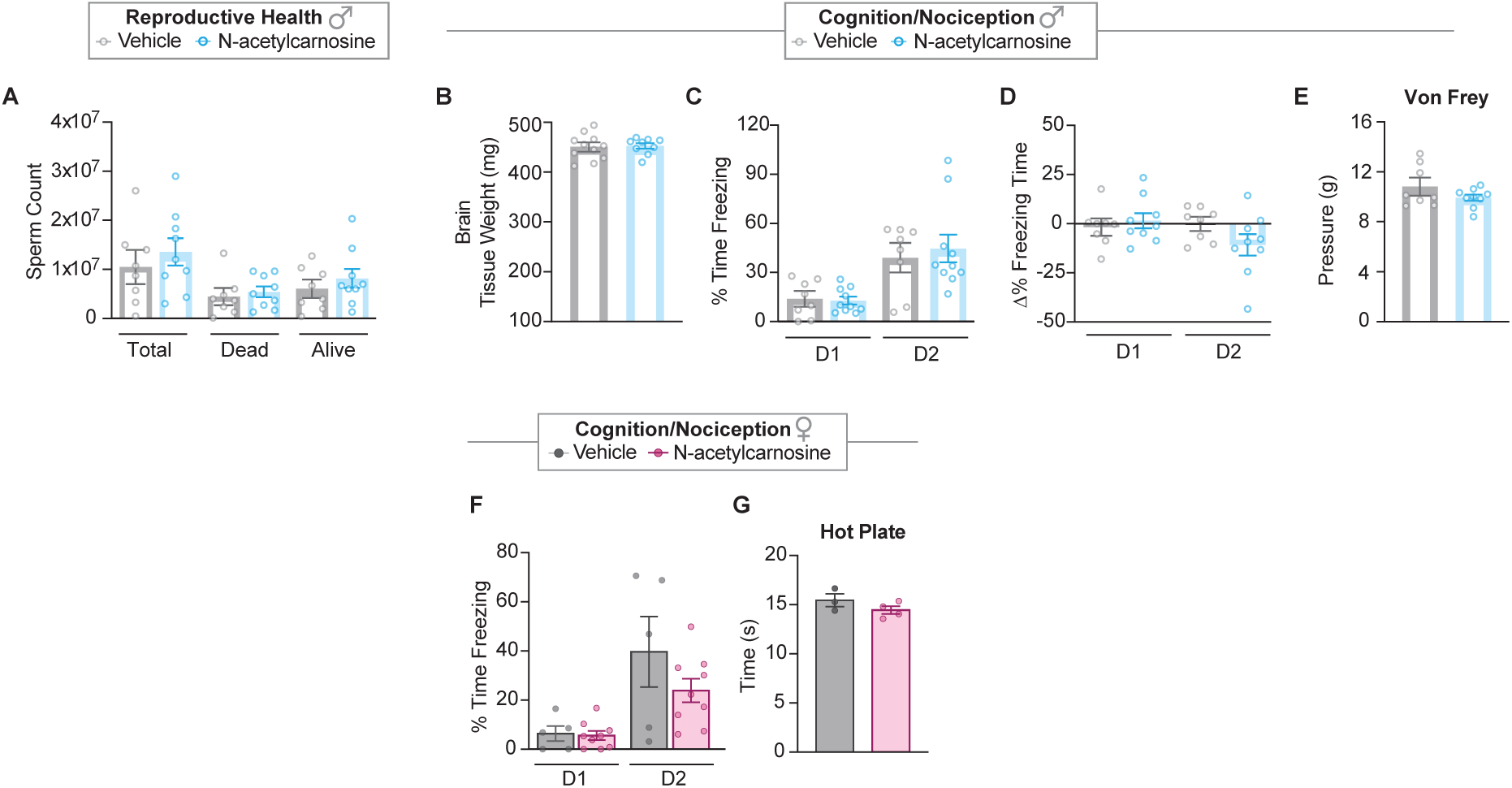
N-acetylcarnosine supplementation does not improve male reproductive health or male and female neurophysiology. Sperm collected from the epididymis of male mice were collected and a sperm count was performed at the time of euthanasia. **A)** N-acetylcarnosine did not affect total, dead, or alive sperm counts. **B)** Brain weights, and cognition as assessed by contextual and conditional fear conditioning **(C and D)** were not affected by N-acetylcarnosine in male mice. **E)** Peripheral pain perception as assess by Von Frey assessment were not affected by N-acetylcarnosine treatment in male mice. Like male mice, contextual fear conditioning **(F)** was not affected by N-acetylcarnosine. **G)** Hot plate assay like Von Frey indicated no effect of N-acetylcarnosine in peripheral nociception in female mice. Data are presented as mean ± SEM and assessed by ttest or 2-way RMANOVA with Bonferroni correction applied for post hoc testing where appropriate. Comparisons were considered statistically significant if *p*<0.05.

**Supplemental Figure 5.**
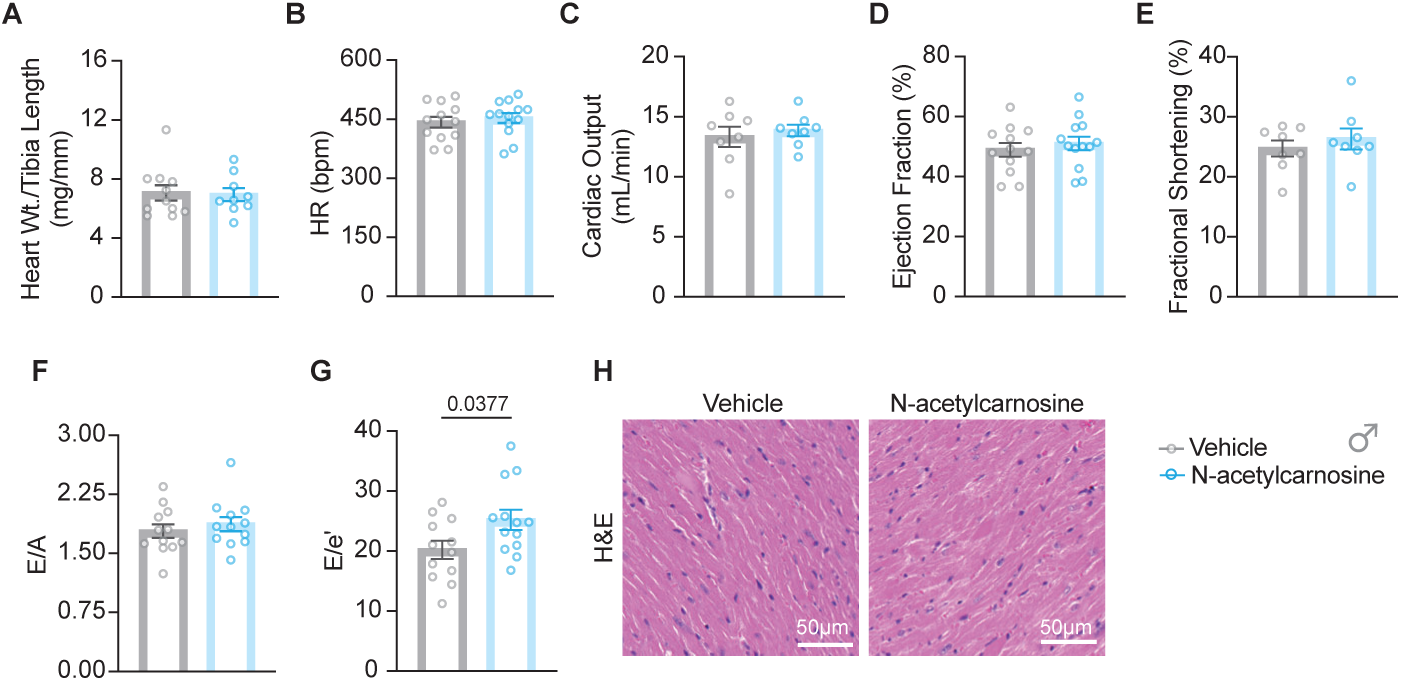
N-acetylcarnosine supplementation improves diastolic function in male mice. **A)** Heart weight relative to tibia length was not different between groups in male mice. Echocardiographic assessment (n=12 vehicle, n=13 N-acetylcarnosine) of mouse systolic function **(B-E)** was also not affected by N-acetylcarnosine in male mice. Early stage **(F)** left ventricular filling was also not different but late diastolic filling **(G)** was greater in N-acetylcarnosine-treated male mice indicating some mild improvements in diastolic function in male mice. **H)** Heart histology as assessed by H&E staining was morphologically similar between groups. Data are presented as mean ± SEM and assessed by unpaired t-test whereby comparisons were considered statistically significant if *p*<0.05.

**Supplemental Figure 6.**
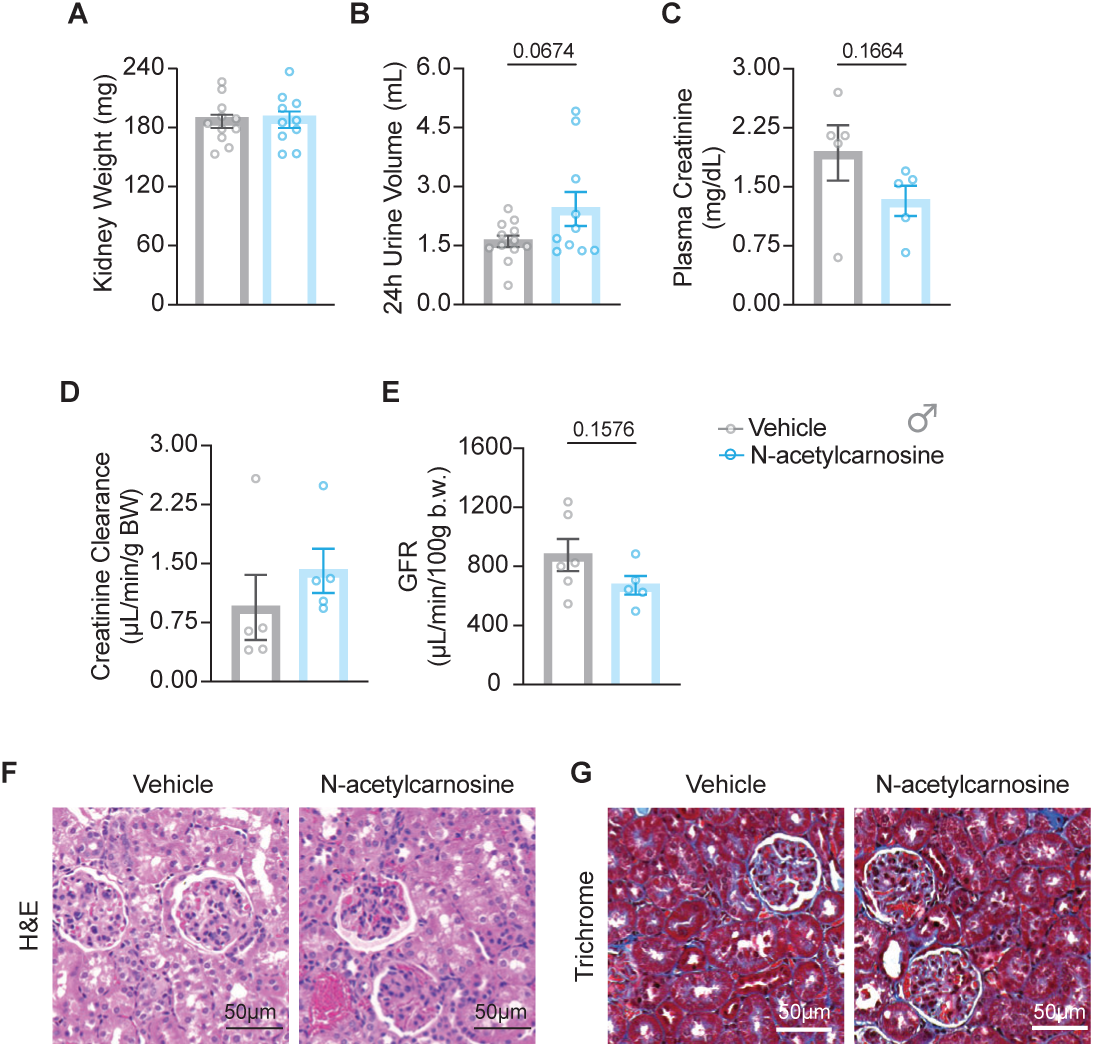
N-acetylcarnosine supplementation does not affect renal function in male mice. **A)** Kidney weight relative to body mass was not different in male mice. 24-hour urine collection was conducted in a modified single mouse cage. **B)** Urine volume trended higher but was not significant in N-acetylcarnosine-treated male mice whereas plasma creatinine concentrations **(C)** trended lower and creatinine clearance **(D)** was equivocal between groups in male mice. **E)** GFR was determined in a subset of male mice (n=6 vehicle, n=6 N-acetylcarnosine) by measuring the excretion of FIT-c labeled senestrin into the urine and was not different between groups. Kidney H&E **(F)** and trichrome **(G)** did not reveal obvious differences in kidney morphology or fibrosis between groups. Data are presented as mean ± SEM and assessed by unpaired t-test whereby comparisons were considered statistically significant if *p*<0.05.

**Supplemental Figure 7.**
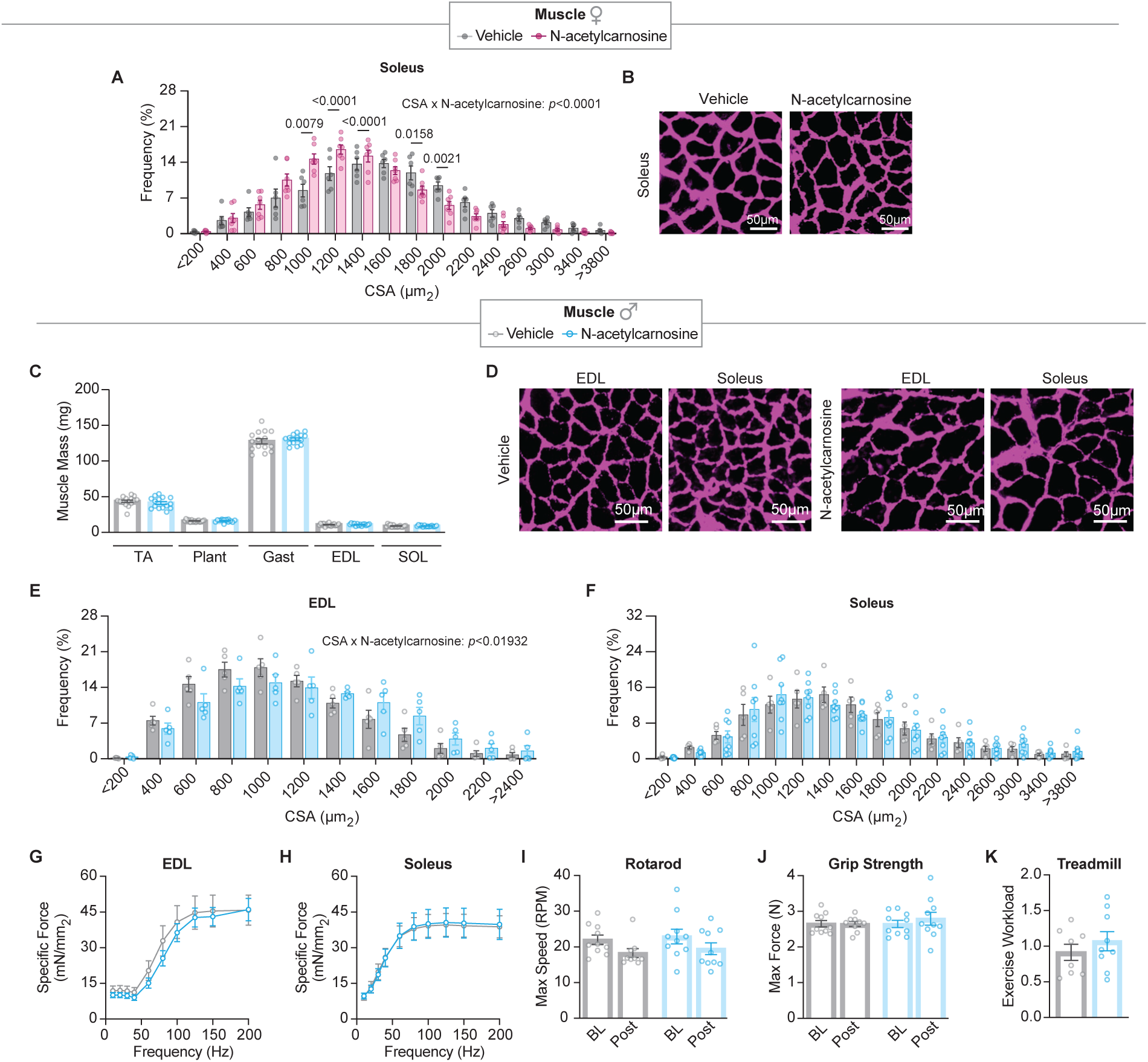
N-Acetylcarnosine supplementation does not dramatically alter muscle function in male mice. **A)** CSA frequency and **(B)** representative image of soleus cross section from female mice demonstrate the frequency of soleus muscle fibers trended smaller in N-acetylcarnosine treated female mice (n=7 vehicle, n=6 N-acetylcarnosine). **C)** Hindlimb muscle weights are not affected by N-acetylcarnosine in male mice. **D)** Representative image of soleus and EDL muscles cryosectioned and stained for laminin for determination of muscle fiber cross sectional area (CSA). **E)** Frequency of larger fibers is increased in EDL muscles from N-acetylcarnosine treated male mice whereas no such effect was found in soleus **(F)**. *Ex vivo* specific force assessed in EDL **(G)** and soleus **(H)** muscles were also not different between groups in male mice. Despite greater CSA in EDL, rotarod **(I)**, grip strength of all four limbs **(J)**, and treadmill work capacity **(K)** were not affected by N-acetylcarnosine treatment. Data are presented as mean ± SEM and assessed by t-test or two-way repeated-measures ANOVA with Bonferroni correction applied for post hoc testing where appropriate. Comparisons were considered statistically significant if *p*<0.05.

**Supplemental Figure 8.**
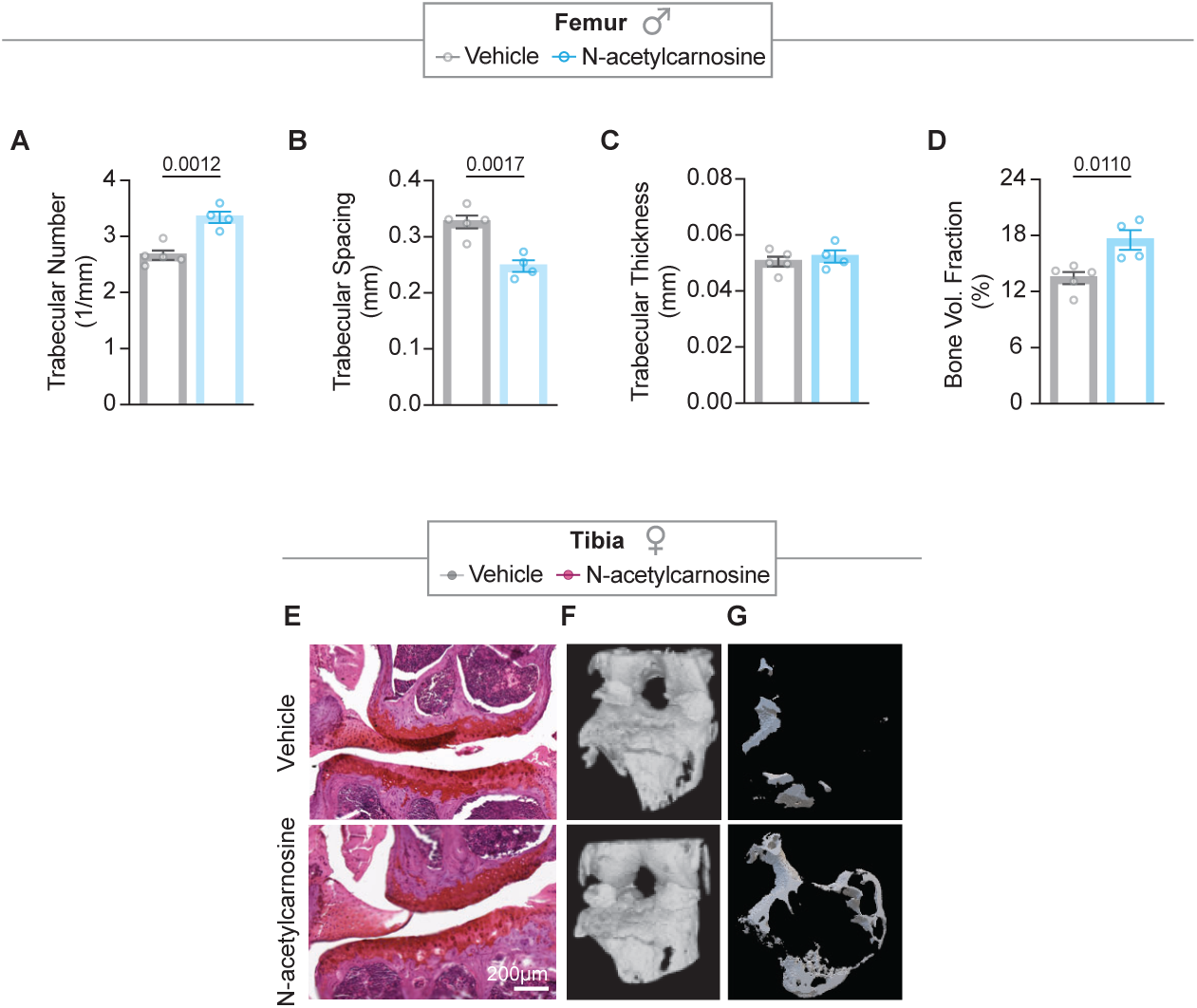
N-acetylcarnosine supplementation improves bone density in femurs from male mice but does not affect knee joint inflammation or tibia bone density. Micro-CT (µCT) analysis of femur bones demonstrated increased trabecular number **(A)**, decreased trabecular spacing **(B)**, no change in trabecular thickness **(C)**, and increased bone volume fraction **(D)** collectively indicating greater bone density in male N-acetylcarnosine treated mice. **E)** Representative images of safranin-O-stained knee joints subjected to Mankin scoring (n=10 vehicle, n=17 N-acetylcarnosine, not significant, data not shown). Representative µCT images of the whole knee joint **(F)** and the tibial epiphysis **(G)**. Trabecular number, spacing, thickness, and bone volume were not different between groups by µCT. Data are presented as mean ± SEM and assessed by t-test or two-way repeated-measures ANOVA with Bonferroni correction applied for post hoc testing where appropriate. Comparisons were considered statistically significant if *p*<0.05.

## Notes

### Competing Interest Statement

The authors have declared no competing interest.

### Summary of Updates

A new data added and figures rearranged for better clarity.

